# Comparison of extraction methods for intracellular metabolomics

**DOI:** 10.1101/2021.12.15.470649

**Authors:** Carolin Andresen, Tobias Boch, Hagen M. Gegner, Nils Mechtel, Andreas Narr, Emrullah Birgin, Erik Rasbach, Nuh Rahbari, Andreas Trumpp, Gernot Poschet, Daniel Hübschmann

## Abstract

Measurements of metabolic compounds inside cells or tissues are of high informative potential since they represent the endpoint of biological information flow and a snapshot of the integration of many regulatory processes. However, it requires careful extraction to quantify their abundance. Here we present a comprehensive study using ten extraction protocols on four human sample types (liver tissue, bone marrow, HL60 and HEK cells) targeting 630 metabolites of different chemical classes. We show that the extraction efficiency and stability is highly variable across protocols and tissues by using different quality metrics including the limit of detection and variability between replicates as well as the sum of concentration as a global estimate of extraction stability. The profile of extracted metabolites depends on the used solvents - an observation which has implications for measurements of different sample types and metabolic compounds of interest. To identify the optimal extraction method for future metabolomics studies, the benchmark dataset was implemented in an easy-to-use, interactive and flexible online resource (R/shiny app MetaboExtract).

## Introduction

Recent developments in high throughput technologies have enabled precise characterization of biological specimens and insight into health and disease. Representing the biological endpoint of the omics cascade, metabolomics is of particular interest and importance with manifold applications, including investigation of neurometabolic disease^1–3^, insights into infectious disease^4–6^ and the microbiome^7,8^ or cancer^9,10^.

Technologies for metabolic measurements, which almost exclusively rely on mass spectrometry coupled to various other techniques, e.g., gas or liquid chromatography, can be coarsely grouped into untargeted and targeted approaches. While untargeted analyses of the metabolome are particularly well suited for exploratory projects and hypothesis generation, they most often do not allow absolute quantification of the different metabolites and many altered features cannot be identified via public database searches. Some technologies from the spectrum of targeted metabolomic analyses allow for absolute quantification. Using combinations of internal standards and calibrants, the LC-MS/MS based Biocrates MxP® Quant 500 kit is an example of such a technology, measuring up to 630 different metabolites from various metabolite classes and providing absolute quantification for a subset of these metabolites, depending on the respective limits of detection (LOD) and the concentrations in the investigated sample type.

Metabolic measurements, especially for absolute quantification, are well established for the analysis of body fluids (blood plasma^11^, cerebrospinal fluid (CSF)^12^, or urine^13,14^). However, little standardization is achieved and no consensus is established for intracellular measurements of large numbers of metabolites belonging to different classes with diverse chemical properties - yet the metabolic processes inside the cell as an atomic unit of life are highly informative. Measuring the intracellular metabolome requires stringent standard operating procedures (SOPs) to limit any variance that may be introduced during this pre-analytical phase. It includes the disintegration of the 3D tissue architecture and the cellular context, the removal of as many extracellular compounds as possible, and the lysis of cells. These steps are followed by extraction protocols which are essential for effective quantification of intracellular abundance of metabolites with heterogeneous chemical characteristics.

In this work, we compare ten different extraction protocols (among which one had two different resolving volumes) for intracellular metabolomic measurements with the Biocrates MxP® Quant 500 kit from four different human sample types, among which we investigated tissues (liver and bone marrow) and cell lines (adherent: HEK (“human embryonic kidney” cell line) and non-adherent: HL60 (“human leukemia 60” cell line)). We further debut our Shiny app “MetaboExtract” to thoroughly explore this comprehensive dataset, enabling in-depth visualization and analysis of the measured metabolites for a given extraction protocol and sample type.

## Methods & Materials

### Samples

Four different sample types were analyzed within this study representing cell culture and primary sample conditions: non-adherent HL60 cells, adherent HEK cells, primary human liver cells and primary human total bone marrow cells. All primary human material in this study was obtained and used following institutional review board approval by the Medical Ethics Committee II of the Medical Faculty Mannheim, University of Heidelberg, Germany in accordance with the declaration of Helsinki after informed written consent.

#### Human bone marrow sample preparation

Bone marrow aspiration from a 39-year-old healthy male volunteer was performed according to standard clinical protocols. Mononuclear cells (MNCs) were isolated from fresh bone marrow (BM) by ficoll density gradient centrifugation. To do so, BM was diluted 1:2 with phosphate-buffered saline (PBS) and loaded on top of ficoll without disturbing the layer. Samples were centrifuged at 400 x g at room temperature for 30 min. MNCs were extracted and washed with PBS twice. Finally, 3×10^6^ cells per sample were collected and snap frozen using liquid nitrogen.

#### HL-60 and HEK sample preparation

HL60 cells were kept under cell culture conditions in DMEM Glutamax with 10% FCS and 1% Penicillin/Streptomycin. HEK cells were kept under cell culture conditions in DMEM Glutamax with 10% FCS and 1% Penicillin/Streptomycin. Cells were washed twice with ice-cold PBS and aliquots of 3×10^6^ cells were snap frozen using liquid nitrogen.

#### Human liver sample preparation

The liver sample was obtained during surgical liver resection of a 75-year-old male patient with hepatocellular carcinoma. Immediately after surgical resection a piece of healthy liver tissue was washed with ice cold 0.9% NaCl solution and snap frozen in liquid nitrogen. Tissue was pulverized to a fine powder without defrosting using a ball-mill (2x 30 sec; 30 Hz; MM 400; Retsch) and pre-cooled stainless-steel beakers. Until extraction, all human samples were stored at -80 °C in 30 mg aliquots.

### Extraction

The ten different extraction protocols compared in this study are described in Figure 1. The protocol 75 EtOH/MTBE was applied with two resolving volumes A) 120 μl and B) 60 μl.

**Figure 1.**
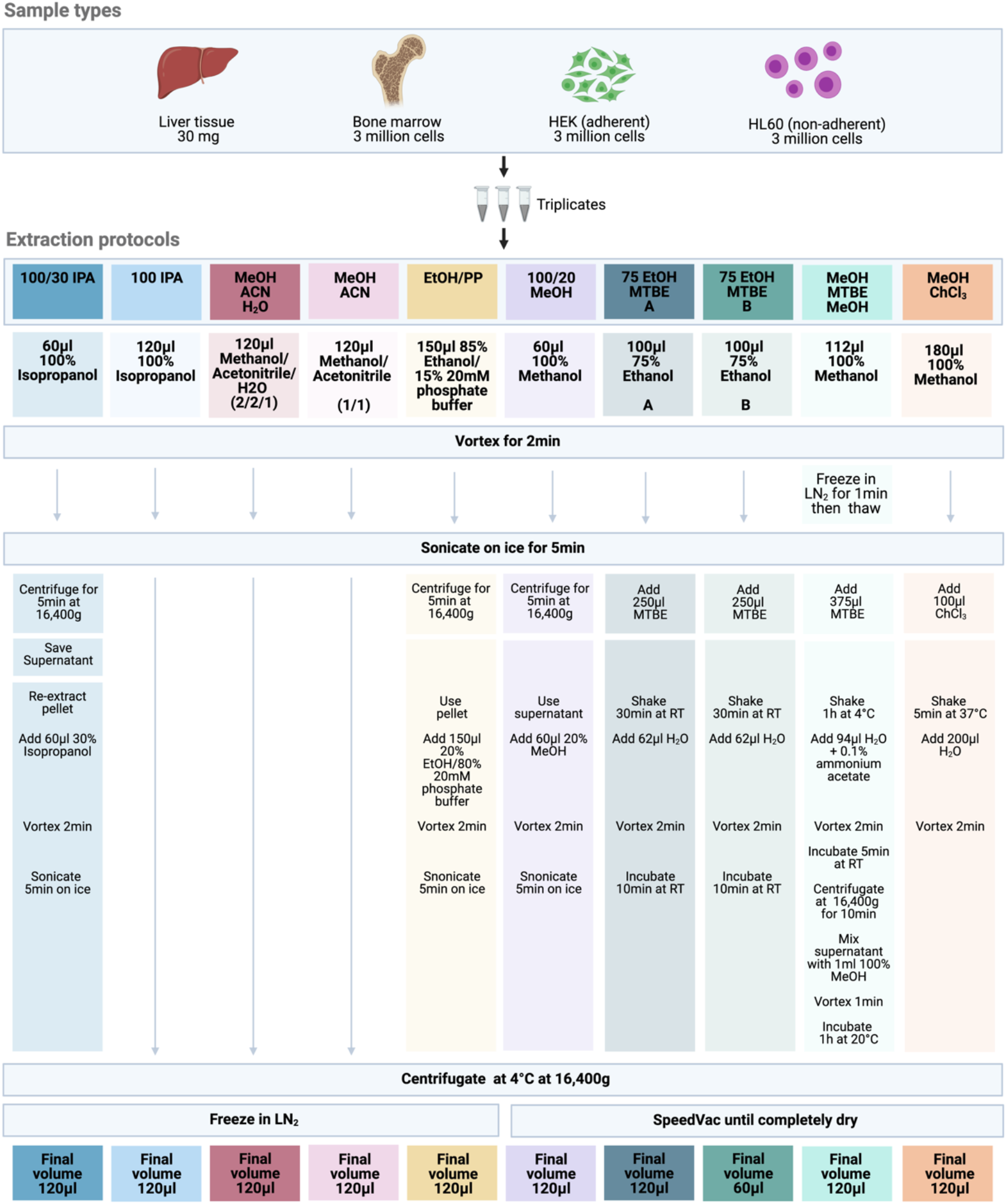
Experimental design and extraction protocol overview. Abbreviations: IPA: isopropanol; MTBE: methyl tert-butyl ether; RT: room temperature

### Sample analysis

In total, 630 metabolites covering 14 small molecule and 9 different lipid classes were analyzed using the MxP® Quant 500 kit (Biocrates) following the manufacturer’s protocol. For a full list of metabolites covered, we refer to the specification of the Biocrates MxP® Quant 500 kit [https://biocrates.com/wp-content/uploads/2021/01/biocrates-Quant500-list-of-metabolites-v4-2021.pdf] (Supplementary Table 1). Of note, 12 different lipid classes are summarized into 9 by the MetIDQ software (Biocrates) merging lysophosphatidylcholines and phosphatidylcholines to glycerophospholipids as well as hexosyl-, dihexosyl- and trihexosylceramides to glycosylceramides. In brief, 10 μl of reconstituted samples were pipetted on a 96 well-plate containing internal standards and dried under a nitrogen stream using a positive pressure manifold (Waters). 50 μl of a 5% phenyl isothiocyanate (PITC) solution were added to each well to derivatize amino acids and biogenic amines. After 1 h incubation time at room temperature, the plate was dried again. To extract the metabolites 300 μl 5 mM ammonium acetate in methanol were pipetted to each filter and incubated for 30 min. The extract was eluted into a new 96-well plate using positive pressure. For further LC-MS/MS analyses 150 μl of the extract was diluted with an equal volume of ultra-pure water. For FIA-MS/MS analyses 10 μl extract was diluted with 490 μl of FIA solvent (provided by Biocrates). After dilution, LC-MS/MS and FIA-MS/MS measurements were performed. For chromatographic separation an UPLC I-class PLUS (Waters) system was used coupled to a SCIEX QTRAP 6500+ mass spectrometry system in electrospray ionization (ESI) mode. Data was recorded using the Analyst (Sciex) software suite and transferred to the MetIDQ software (version Oxygen-DB110-3005) which was used for further data processing, i.e. technical validation, quantification and data export. All metabolites were identified using isotopically labeled internal standards and multiple reaction monitoring (MRM) using optimized MS conditions as provided by Biocrates. For quantification either a seven-point calibration curve or one point calibration was used depending on the metabolite class.

### Data analysis

Downstream analysis of MetIDQ output was performed using custom R scripts (v4.0.0). The code used for the figures in this paper is available at https://github.com/andresenc/extractioncomparison. ggplot2 (v3.3.3) and cowplot (v1.1.1) were used for generation of plots.

Metabolites were defined as ‘above LOD’ when at least 2 replicates met this criterion. For PCA, raw data was filtered for metabolites that were below LOD in all samples, zero values were replaced by taking the minimum of all measured concentrations per metabolite and adding 20% to this value, data were log2-transformed and scaled using the Pareto method as described by van den Berg et al.^15^. To identify associations between principal components and experimental setting, the Kruskal-Wallis test was applied. P-values were corrected using Benjamini-Hochberg and considered significant if p.adjust < 0.05.

To compare concentration yields between extraction protocols, an ANOVA was calculated based on log2-transformed data. The ten extraction protocols were used as the categorical variable and concentration as the dependent variable. Using the Tukey post-hoc test group labeling, the optimal extraction methods with the highest median yield were determined for each metabolite and sample type.

### Data availability

Data can be explored and downloaded using the Shiny app MetaboExtract which is available at http://www.metaboextract.shiny.dkfz.de. The underlying code is also available at https://github.com/andresenc/MetaboExtract.

## Results

In order to perform measurements of intracellular metabolite concentrations, cells or tissue need to be lysed and metabolites have to be extracted. In this work, we compared ten different extraction protocols on four sample types and assessed their effect on abundance and reproducibility of the determined metabolite concentrations and on the number of metabolites for which concentrations above the limit of detection (LOD) could be obtained (Figure 1). The MxP® Quant 500 kit covers up to 630 metabolites from different chemical classes. These classes with their respective numbers of metabolites are displayed in Figure 2a, highlighting the large fraction of triacylglycerols (242/630 = 38.4%) and glycerophospholipids (90/630 = 14.3%).

**Figure 2.**
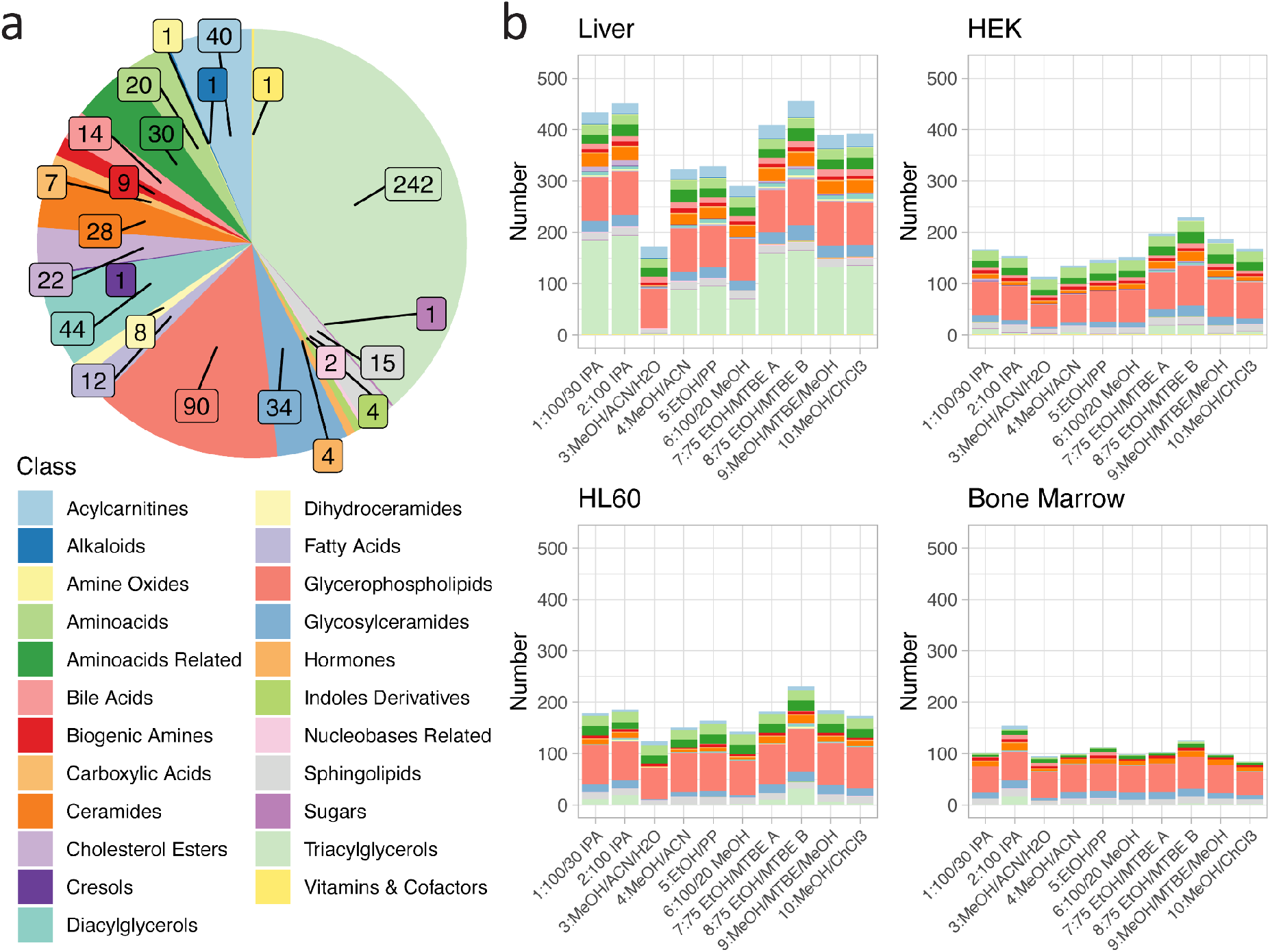
(a) Metabolites in the Biocrates MxP® Quant 500 kit. Colours encode the different metabolite classes. Numbers of metabolites per class are indicated by inset. (b) Metabolites above limit of detection (LOD) in the four different sample types and for all extraction protocols. Colours encode metabolite classes as in (a).

### Effective quantification depends on sample type and extraction protocol

Effective quantification of metabolites requires a reliable signal. The LOD is defined as 3 times signal-to-noise of the baseline, calculated by the software MetIDQ for each metabolite. Figure 2b displays the numbers of metabolites above LOD, colour coded by the different metabolite classes, in four different cell types as stacked bar graphs for the different extraction protocols. On average, the highest number of metabolites above LOD was observed for liver tissue (median: 391, range 171 - 456), the lowest number was observed in bone marrow (median: 101, range 85 - 154), while the two investigated cell lines, one adherent (HEK, median: 160.5, range 113 - 230) and one non-adherent (HL60, median: 176, range 124 - 231) had very similar and intermediate total numbers of metabolites above LOD. The distribution of metabolites above LOD among the different metabolite classes was similar to the overall distribution of metabolites in the kit (Figure 2), with the important additional observation that the most variable class was triacylglycerols (median +/- median absolute deviation (MAD): 3.5 +/- 5.2). For the four different sample types, different extraction protocols performed best according to the number of metabolites above the LOD: while for liver tissue and HL60, extraction protocols 8: *75 EtOH/MTBE B* and 2: *100 IPA* performed best, protocols 7: *75 EtOH/MTBE A* and 8: *75 EtOH/MTBE B* performed best for the HEK cell line and 2: *100 IPA* and 8: *75 EtOH/MTBE B* performed best for the bone marrow samples. Integrating across the different tissues, protocol 8: *75 EtOH/MTBE B* performed best. Figure 3a also displays the number of metabolites above LOD but aggregated and dodged by extraction protocol. The one method which performed worst across all sample types was 3: *MeOH/ACN/H2O*, and all methods containing methanol had comparably lower yield.

**Figure 3.**
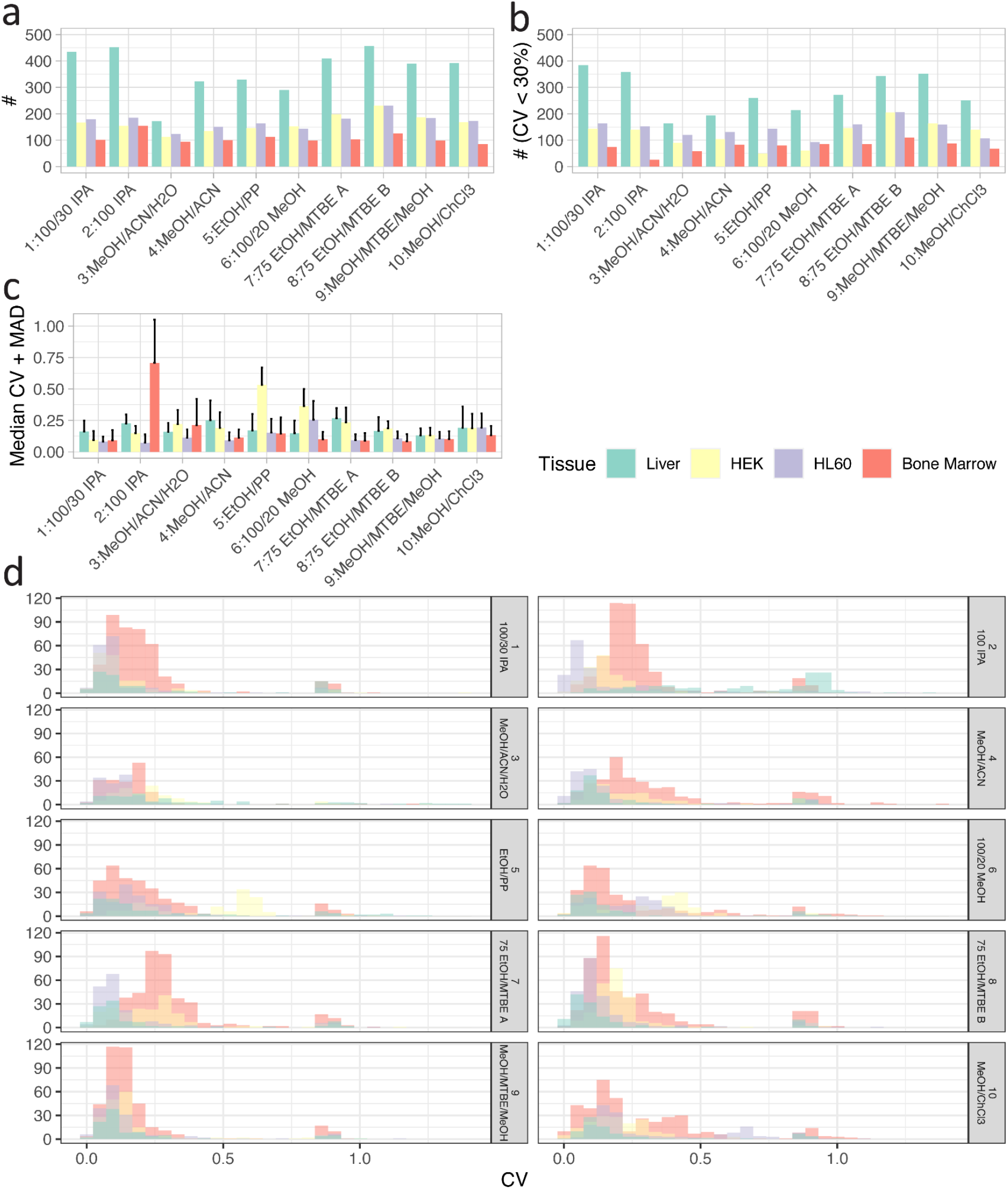
(a) Summary statistics of numbers of metabolites. (b) Numbers of metabolites with coefficients of variation (CV) below 30%. (c) Median and median absolute deviation (MAD) for the CVs for all four cell types and all nine extraction protocols. (d) Distributions of CVs displayed as histograms for different tissues and extraction methods (only metabolites above LOD). Colour code applies to all items.

Furthermore, the metabolite concentrations were assessed. For each metabolite, best extraction methods were determined by running an ANOVA on the measured concentration above the LOD. Methods with either the highest median yield or non-significantly lower concentrations were considered optimal. When ranking extraction methods using this quality metric, the best extraction methods for each tissue were discovered to be the same ones as those found using numbers of metabolites above LOD; heterogeneity was observed only in the lower ranking extraction methods. This again confirmed the importance of a suitable extraction protocol (Supplementary Figure 1).

### Reproducibility of extraction efficacies between replicates

We next assessed the reproducibility of measurements. To this end, we measured biological replicates for all combinations and computed the coefficients of variation (CVs). Figure 3b displays the number of metabolites with CV < 30%, reflecting those metabolites with good reproducibility of extraction efficacy. In agreement with the observation that liver tissue had the highest number of metabolites above LOD in total, liver tissue was also the cell type with the highest number of reproducible metabolite concentrations. In general, the distribution of the numbers of reproducible metabolites across the different extraction protocols correlated well with the total numbers of metabolites above LOD (Figure 3a,b). Of note, for some tissues, a large fraction of all metabolites was reproducible (i.e., had low CVs) in all extraction protocols, e.g., for liver tissue, while for other tissues, only small fractions of all metabolites above LOD were reproducible, especially for bone marrow. This was reflected by the overall distribution of the CVs, median and MAD of which are shown in Figure 3c, with many combinations of cell type and extraction protocol reaching median CVs as low as 20%, but also exhibiting marked outliers, like bone marrow in extraction protocol 2: *100 IPA* and 9: *MeOH/ACN/H20*, HEK in extraction protocol 5: *EtOH/PP* or HL60 in extraction protocol 6: *100/20 MeOH*. Figure 3d shows the distributions of CVs as histograms for the metabolites above LOD. It can be observed that for all combinations of cell type and extraction protocol, the distributions of CVs had a long tail towards high CVs, representing small numbers of metabolites with low reproducibility. Matching the observations described above for Figure 3c, extraction protocol 2: *100 IPA* applied to bone marrow led to a particularly high number of such metabolites. In analogy to Figure 3b, Supplementary Figure 2 displays the histograms of CVs for all metabolites (including those below LOD). As expected, adding metabolites below LOD populated quantiles of metabolites with high CVs.

### Sum of concentrations as a quality measure

In search of useful and intuitive quality control (QC) metrics, the sum over all metabolite concentrations was calculated to estimate the overall analytical reproducibility between replicates. The sum of concentrations (SOC) reflects the variability between replicates, which may indicate the reproducibility of multi-step experimental extraction protocols as well as an impression of the overall extraction efficiency between sample types. Supplementary Figure 3 displays this value for three biological replicates for the four cell types and all extraction protocols. In general, a good consistency over the biological replicates can be observed which depends on the protocol and the tissues. For example, 10: *MeOH/ChCl3* shows low variation for liver and bone marrow (CV: 0.03 and 0.05) but high variation for HEK and HL60 (CV: 0.18 and 0.44). Even though among all extraction protocols 2: *100 IPA* performed best based on the number of metabolites above LOD, the SOC reaches similar values for methods 1, 3, 4, 5, 6, and 8 (c.f. Figure 2b). Furthermore, SOC is a metric which is complementary to the number of metabolites above LOD as it aggregates absolute concentrations and thereby makes use of the functionality of the MxP® Quant 500 kit. Due to the different nature of samples, concentrations for HEK, HL60 and bone marrow are given in picomole per 10^6^ cells and in picomole per mg for liver tissue. Therefore, the SOC for the liver is displayed on a different scale and the data shown here does not contradict the observation that liver tissue showed the highest number of metabolites above the LOD. However, using published values for hepatocellularity of (65 – 185) · 10^6^ cells/g^16^ or (139 ± 25) · 10^6^ cells/g^17^, the 30 mg of liver tissue correspond to (1.95 – 5.55) · 10^6^ cells or (4.17 ± 0.75) · 10^6^ cells, respectively, and are in a similar range as the 3 · 10^6^ cells of the other sample types. For HEK, HL60 and bone marrow, there is an overall trend of the SOC from high to low for the sample types respectively. A comparison of the SOC with the number of metabolites above the LOD as a QC metric shows that these two measures do not correlate and that the SOC can provide additional information (e.g. 2: *100 IPA* for HEK and HL60, c.f. Figure 2b and Supplementary Figure 3).

### Metabolite concentrations

Restricting our attention to those metabolites whose concentrations are above LOD and which have acceptable reproducibility, we compiled absolute concentrations of these metabolites. Figure 4a shows extraction profiles of these absolute concentrations for all cell types obtained with extraction protocol 8: *75 EtOH/MTBE B*, and Figure 4b shows extraction profiles for all remaining extraction protocols for liver tissue. The complete information for all cell types and all extraction protocols is shown in Supplementary Figure 4. The concentrations in HEK cells show the broadest range, from 0.015 pmol/10^6^ cells for glycolithocholic acid sulfate (a bile acid), to 68.6 nmol/10^6^ cells for hexose. Even though in liver tissue and HEK cells hexose has the overall highest concentration, in HL60 and bone marrow, the abundance of hexose is below LOD in all extraction methods (Figure 4a and Supplementary Figure 4).

**Figure 4.**
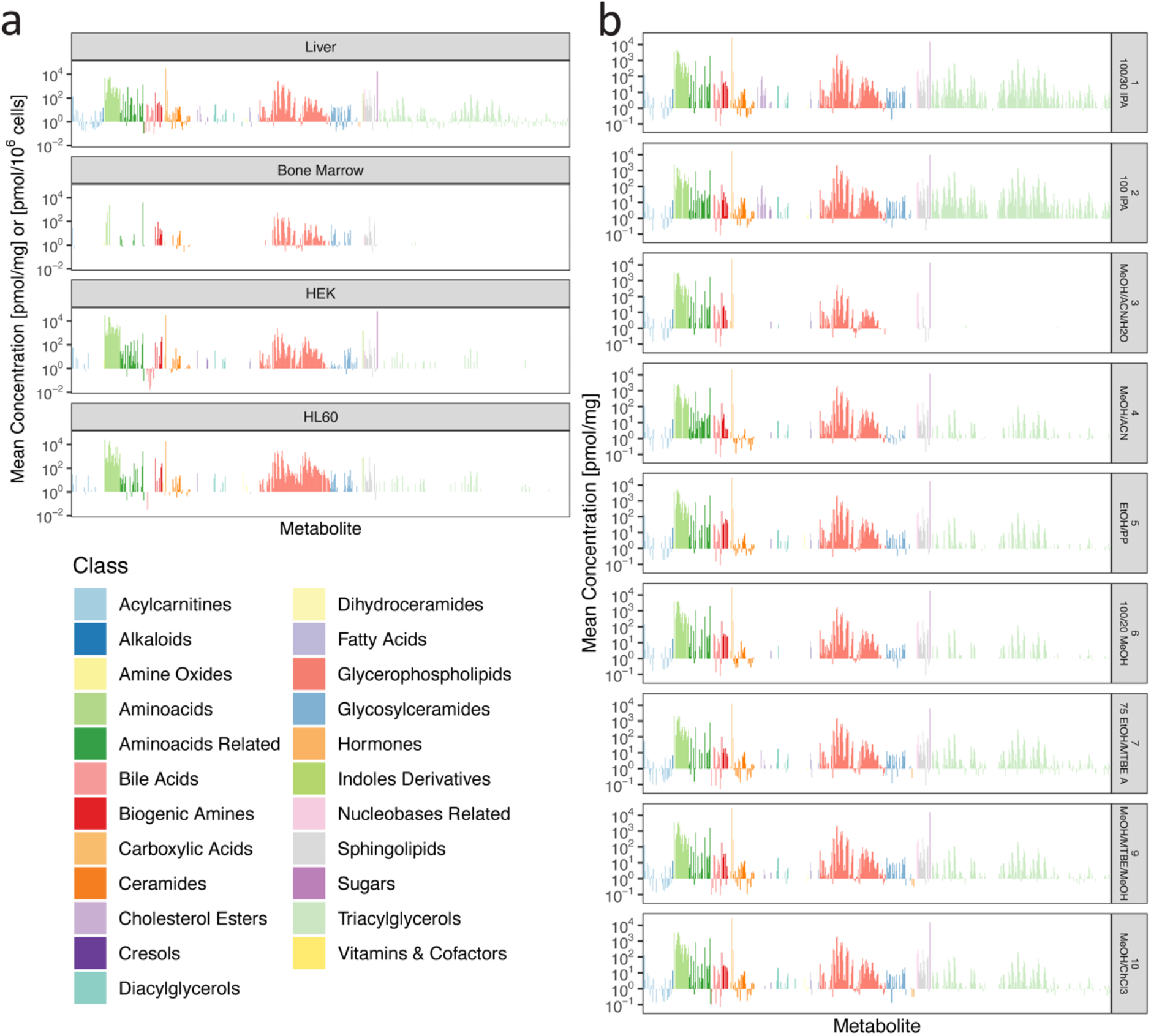
Mean absolute concentrations between replicates. (a) Comparison of all four cell types for 8: *75 EtOH/MTBE B*. Concentrations of bone marrow, HEK and HL60 samples are plotted as pmol/10^6^ cells and pmol/mg for liver tissue. (b) Comparison of all additional extraction protocols for liver tissue. Colours encode metabolite classes as in (a).

### Variance across extraction profiles is driven by sample type and solvent

An unsupervised principal component analysis (PCA) showed that biological replicates clustered together, and that the first five principal components (PCs) had their variance explained mostly by tissue type as the main effect. PC3-6 and PC9-10 reflected extraction protocol (Figure 5a,c). Focusing only on liver tissue, PCA revealed that similar protocols clustered together and the main variability between the extraction methods is determined by the solvents. PC1, which captures the highest fraction of the total variance, is most strongly associated with IPA, with next highest association strengths for MeOH and ACN (Figure 5b,d). For PC1, the highest loadings contain multiple triacylglycerides, whereas ceramides and sphingomyelins seem to drive the variability on PC2. This reflects that extraction efficiencies for the various metabolite classes are highly different across the different solvents, depending on the chemical properties of the metabolite classes (Figure 5e and Supplementary Figure 5).

**Figure 5.**
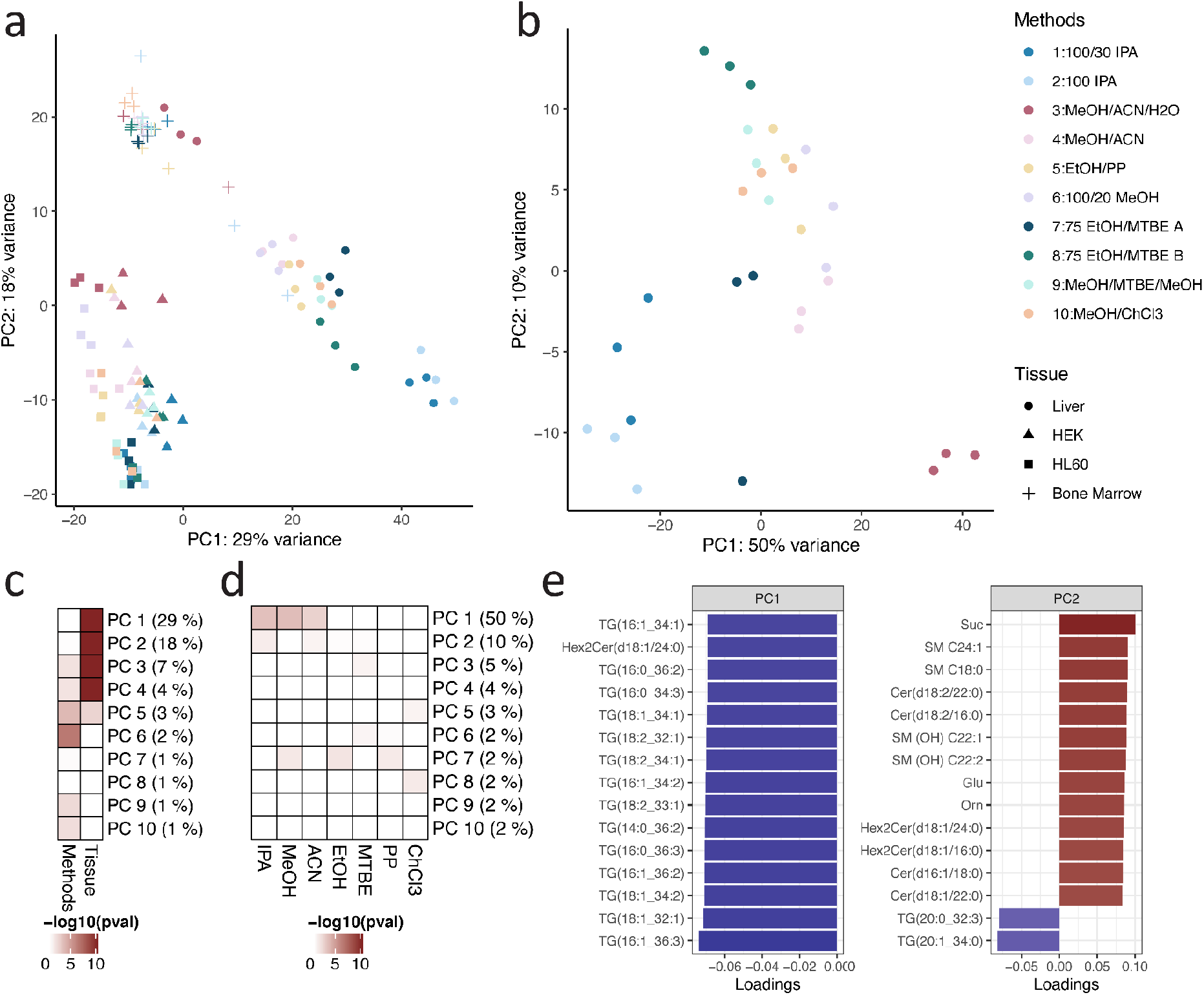
(a) PCA of the ten different extraction protocols used across all sample types. (b) PCA of the ten different extraction protocols used on liver tissue samples. Colours encode metabolite classes as in (b). (c) Heatmap showing association strength between PCs and experimental setting (extraction protocol or tissue), colour coding by p-value of the Kruskal-Wallis test. (d) Heatmap showing association strength between PCs and solvents used in extraction protocols, colour coding by p-value of the Kruskal-Wallis test. (e) Barplot of the 15 highest (ranked by absolute value) loadings for PC1 and PC2. The colour encodes the signs of the loadings.

### A resource for interactive exploration

Efficient and reproducible extraction of metabolites is crucial for a determination of their intracellular abundance. The data presented in Figures 2-4 and in Supplementary Figures 1-4 were made available as an easy-to-use interactive R/shiny app which allows users to explore and compare the different extraction methods at http://www.metaboextract.shiny.dkfz.de. Tissues, methods and classes of metabolites can be (de)selected to focus on the data of interest and the maximal CV between replicates can be chosen to identify the most suitable method for future analyses.

## Discussion

While metabolic measurements are well established for the analysis of body fluids like blood plasma^11^, CSF^12^, or urine^13,14^, there is less consensus on standardization for extraction of intracellular metabolites and subsequent metabolomic measurements. The pre-analytical phase, i.e., the sample collection, handling and pre-treatment in a metabolomic study has tremendous effects on results of the study. While stringent SOPs during sample collection and handling control for this error prone pre-analytical part^18^, the adequate choice of an extraction protocol, i.e., the pre-treatment of the sample to extract the metabolites of interest on a given analytical platform^19,20^ also influences the range, robustness and validity of the generated data. Here, we provide a comprehensive analysis of ten different extraction methods for intracellular metabolic measurement for four human sample types: The tissue types liver and bone marrow, as well as the cell lines HEK (adherent) and HL60 (non-adherent) using the Biocrates MxP® Quant 500 kit, which allows absolute quantification of up to 630 metabolites via LC-MS/MS and FIA-MS/MS measurements.

Assessment of QC metrics (LOD, CV and SOC) showed that efficiency and reproducibility are complementary sources of information. In addition, we used the SOC as a quality metric to assess the global variability between replicates and could show that certain protocols are more prone to technical variability than others. Overall, 2: *100 IPA* and 8: *75 EtOH/MTBE B* showed the best results across the different sample types. However, for each sample type and metabolites of interest, the optimal method may be different.

For liver tissue, 8: *75 EtOH/MTBE B*, 2: *100 IPA* and 1: *100/30 IPA* resulted in the highest, second highest and third highest numbers of metabolites above LOD, respectively, with relatively low differences between these three methods. With more than 400 metabolites above LOD for all these methods, almost the full spectrum of the Biocrates MxP® Quant 500 kit can thus actually be exploited for liver tissue. For bone marrow, 2: *100 IPA*, 8: *75 EtOH/MTBE B* and 5: *EtOH/PP* resulted in the highest, second highest and third highest numbers of metabolites above LOD, respectively, but 2: *100 IPA* performed considerably better than the latter two. With the best methods yielding hardly more than 150 metabolites over LOD, only one quarter of the metabolites measured by the Biocrates MxP® Quant 500 kit are usable. The two different cell lines showed very similar rankings of the different methods, with 8: *75 EtOH/MTBE B* having by far the best yield of metabolites above LOD. With slightly over 200 metabolites above LOD for this method, the extraction worked slightly better than for bone marrow. Of note, the most variable class of metabolites were triacylglycerols, at least partially reflecting variations in content of these metabolites across the different tissues. The one method which performed worst across all sample types was 3: *MeOH/ACN/H2O*, and all methods containing methanol had comparably lower yield. This could potentially be due to the fact that methanol may not be apolar enough to properly extract the very apolar lipid species.

As expected, our data show that the used solvents influence the profile of extracted metabolites due to the different solvent strength and polarity of the organic solvents or mixtures thereof. To ensure quantitative extraction of very hydrophobic lipid species, apolar solvents like isopropanol or MTBE - classical solvents used in lipidomics approaches - are required, while in comparison the influence on extraction efficacy of polar compounds of core carbon and nitrogen metabolism (e.g. amino acids) is minor.

For HEK, HL60 and bone marrow samples, 3×10^6^ cells were used as input, whereas the extraction efficiency was significantly higher for liver samples which had 30 mg tissue as input. This suggests that the extraction efficiency and reproducibility can be increased by larger input numbers.

In addition to this comprehensive overview and comparison of the different extraction protocols that yield generalized conclusions across the sample types, we provide a free and flexible online resource to allow scientists to explore and subset these insights in a tailored manner. Using MetaboExtract, the dataset generated here can be visualized and analyzed depending on specific requirements. It enables further in-depth analysis based on a metabolite of interest, a metabolite class or related to one sample type only. A specific use case could be, e.g., a project which aims at measuring amino acids in HL60. Even though method 3: *MeOH/ACN/H2O* yields the least number of metabolites above LOD in general, for this specific combination of metabolite class and sample type, method 3: *MeOH/ACN/H2O* still may be a good choice.

It has the potential to inform future studies of the most adequate extraction protocol and to optimize the pre-analytical phase in general, i.e., using measurement technologies other than the Biocrates MxP® Quant 500 kit. Lastly, the flexible nature of MetaboExtract has the potential to grow by adding further data sets, e.g., from other tissues or different model organisms, that were generated using this highly standardized assay.

In conclusion, in this work, we provide a comprehensive comparison of different extraction methods for intracellular metabolic measurements by absolute quantification, identify optimal choices of methods for different sample types including primary tissues and cell lines, and provide a free online resource for interactive exploration of the data.

## Supporting information

Supplementary_Items

## Acknowledgement

The authors gratefully acknowledge the data storage service SDS@hd supported by the Ministry of Science, Research and the Arts Baden-Württemberg (MWK), by the German Research Foundation (DFG) through grant INST 35/1314-1 FUGG and INST 35/1503-1 FUGG and by the German Federal Ministry of Education and Research under the funding code 161L0212 – SMART-CARE.

