## Supplementary_Items for "Comparison of extraction methods for intracellular metabolomics"

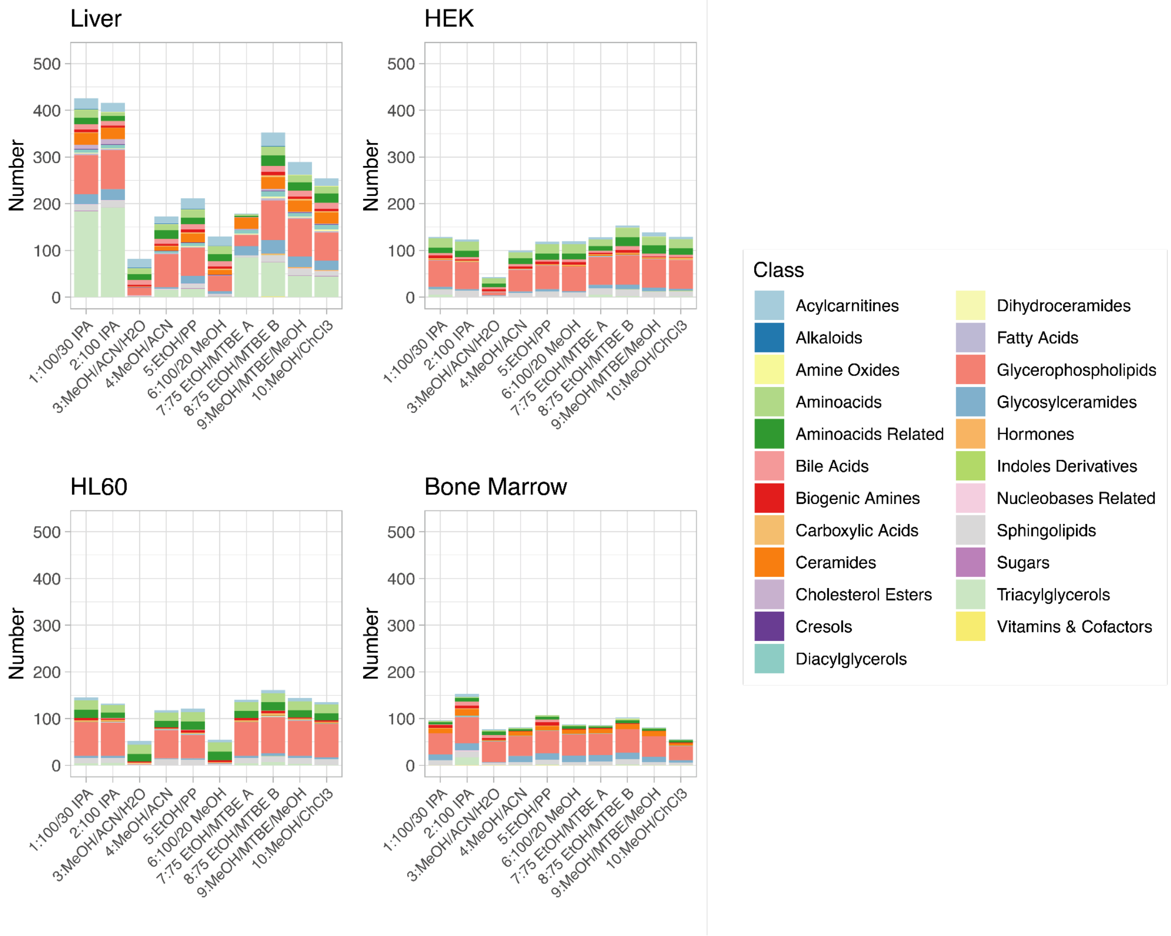


**Suppl. Figure 1:** Metabolites above limit of detection (LOD) with highest concentration yield among extraction protocols in the four different sample types calculated by ANOVA. Colors encode the different metabolite classes.

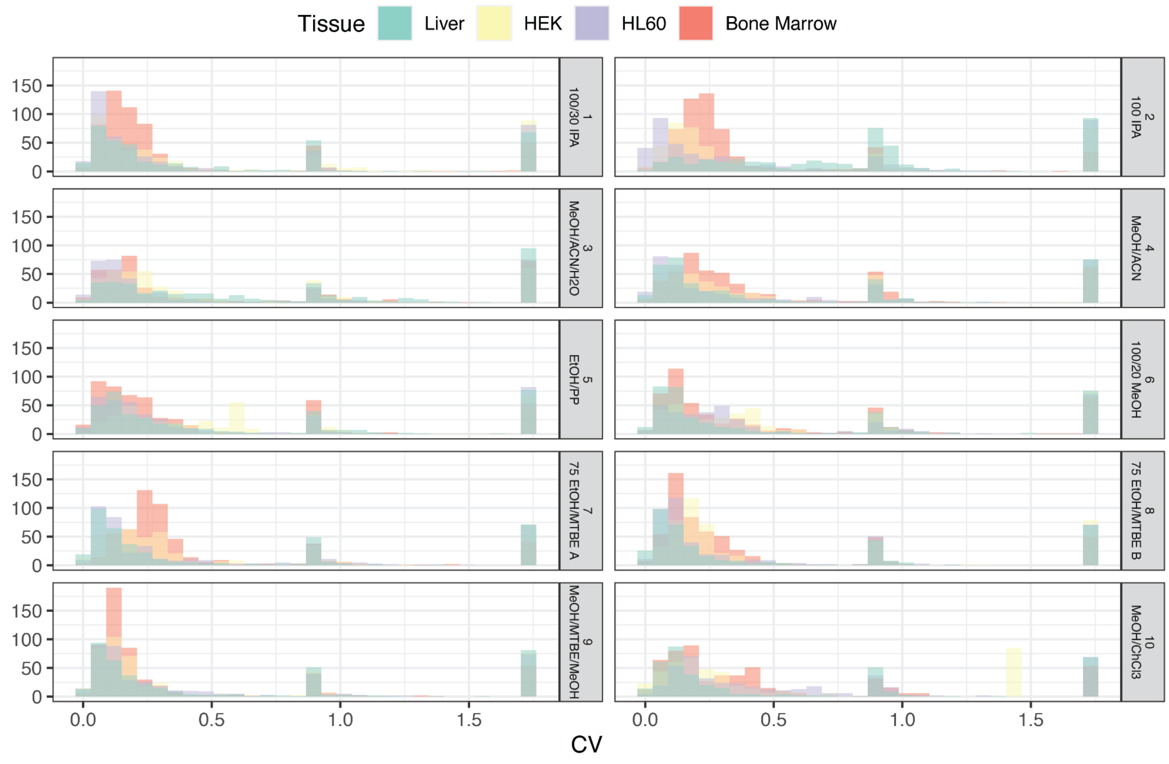


**Suppl. Figure 2:** Distributions of CVs between replicates across extraction protocols and sample types.

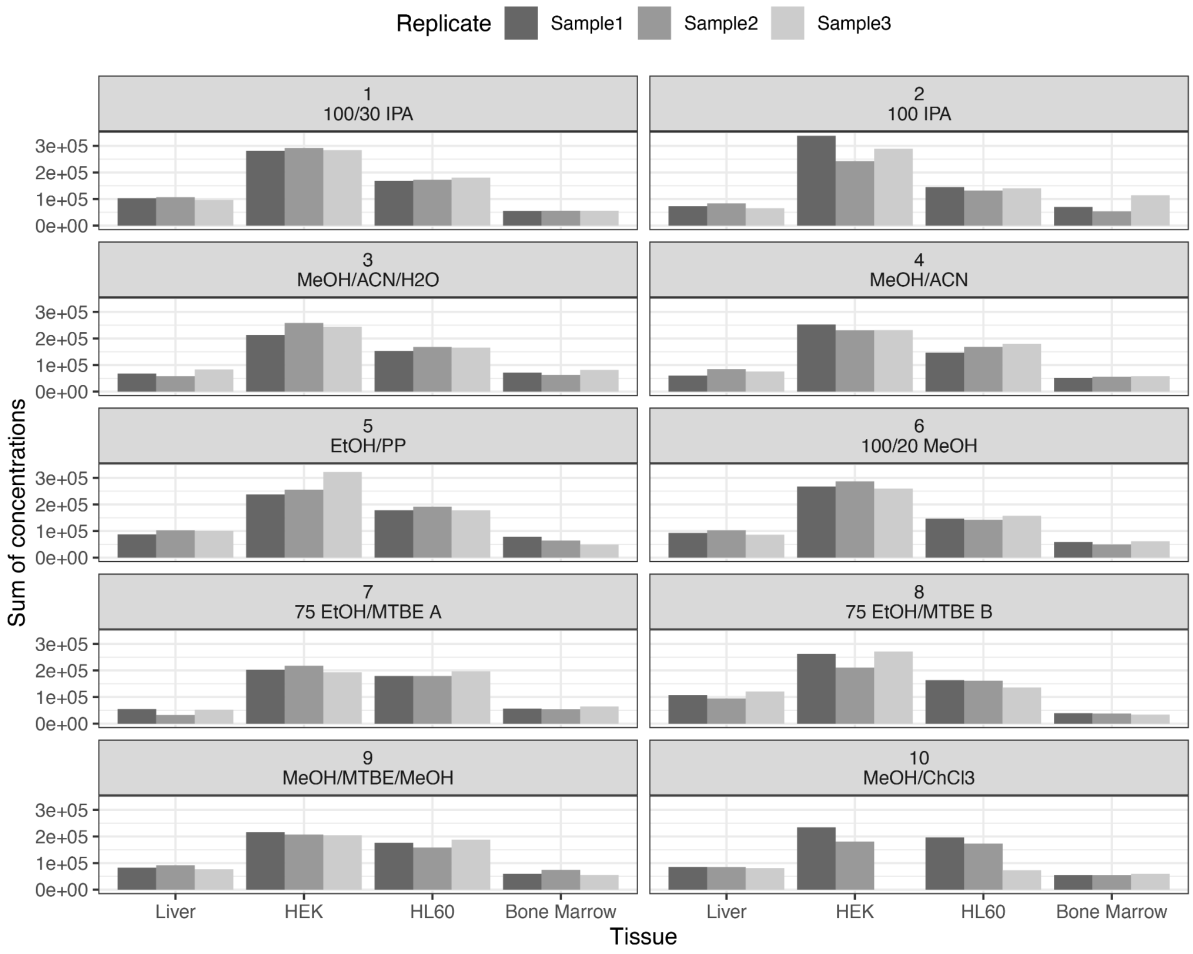


**Suppl. Figure 3:** Sums of concentrations between replicates across extraction protocols and sample types. Of note, one replicate of HEK was removed because of very low concentrations.

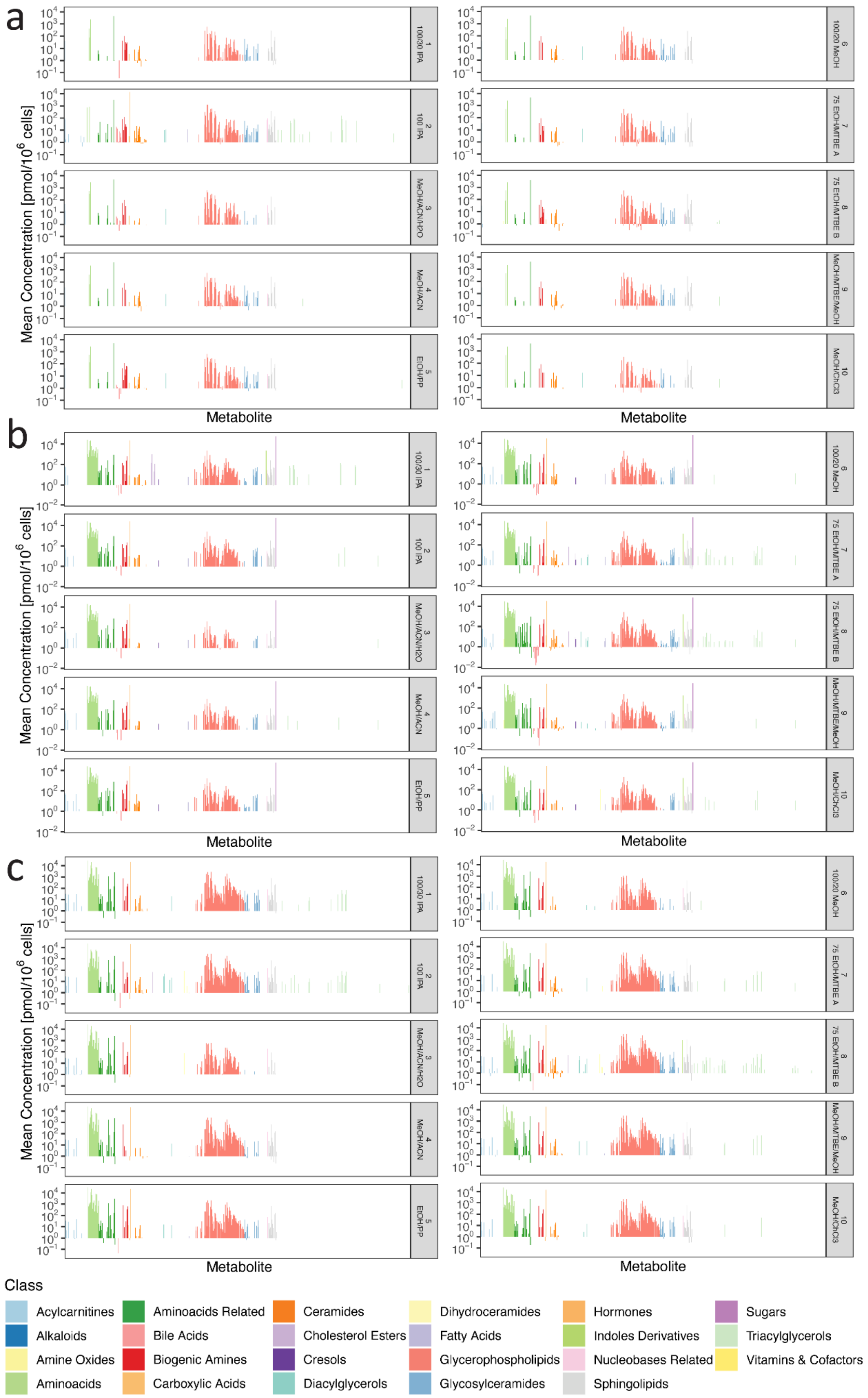


**Suppl. Figure 4:** Mean absolute concentrations between replicates. (a) Bone marrow, (b) HEK and (c) HL60.

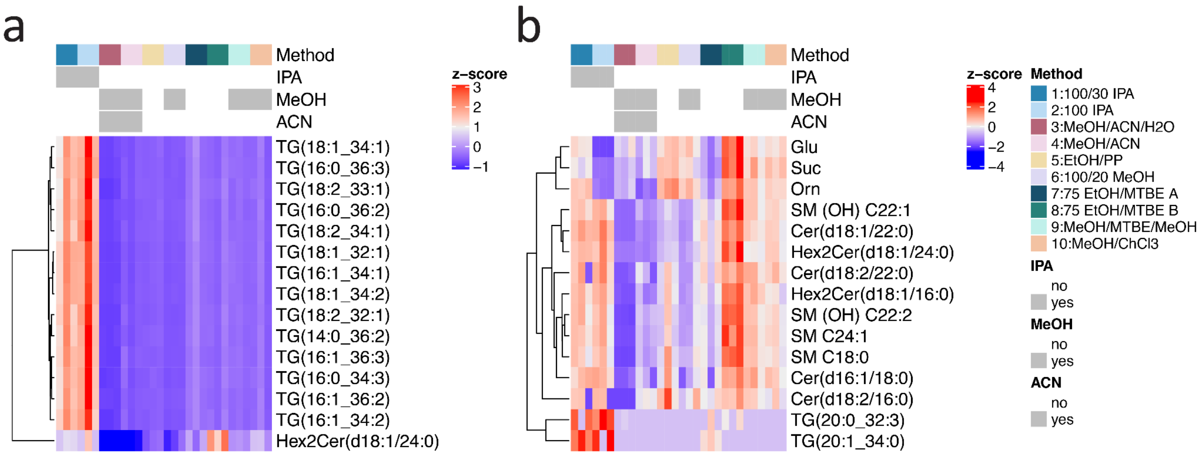


**Suppl. Figure 5:** Concentrations of the 15 highest (ranked by absolute value) loadings (metabolites) for (a) PC1 and (b) PC2. Methods and selected solvents are encoded in (b).

**Suppl. Table 1:** Metabolites in MxP® Quant 500 kit.

| Metabolite | Name | Class |
|---|---|---|
| C0 | Carnitine | Acylcarnitines |
| C2 | Acetylcarnitine | Acylcarnitines |
| C3 | Propionylcarnitine | Acylcarnitines |
| C3-DC (C4-OH) | Malonylcarnitine (Hydroxybutyrylcarnitine) | Acylcarnitines |
| C3-OH | Hydroxypropionylcarnitine | Acylcarnitines |
| C3:1 | Propenylcarnitine | Acylcarnitines |
| C4 | Butyrylcarnitine | Acylcarnitines |
| C4:1 | Butenylcarnitine | Acylcarnitines |
| C5 | Valerylcarnitine | Acylcarnitines |
| C5-DC (C6-OH) | Glutarylcarnitine (Hydroxyhexanoylcarnitine) | Acylcarnitines |
| C5-M-DC | Methylglutarylcarnitine | Acylcarnitines |
| C5-OH (C3-DC-M) | Hydroxyvalerylcarnitine (Methylmalonylcarnitine) | Acylcarnitines |
| C5:1 | Tiglylcarnitine | Acylcarnitines |
| C5:1-DC | Glutaconylcarnitine | Acylcarnitines |
| C6 (C4:1-DC) | Hexanoylcarnitine (Fumarylcarnitine) | Acylcarnitines |
| C6:1 | Hexenoylcarnitine | Acylcarnitines |
| C7-DC | Pimelylcarnitine | Acylcarnitines |
| C8 | Octanoylcarnitine | Acylcarnitines |
| C9 | Nonanoylcarnitine | Acylcarnitines |
| C10 | Decanoylcarnitine | Acylcarnitines |
| C10:1 | Decenoylcarnitine | Acylcarnitines |

| | | |
|---|---|---|
| C10:2 | Decadienoylcarnitine | Acylcarnitines |
| C12 | Dodecanoylcarnitine | Acylcarnitines |
| C12-DC | Dodecanedioylcarnitine | Acylcarnitines |
| C12:1 | Dodecenoylcarnitine | Acylcarnitines |
| C14 | Tetradecanoylcarnitine | Acylcarnitines |
| C14:1 | Tetradecenoylcarnitine | Acylcarnitines |
| C14:1-OH | Hydroxytetradecenoylcarnitine | Acylcarnitines |
| C14:2 | Tetradecadienylcarnitine | Acylcarnitines |
| C14:2-OH | Hydroxytetradecadienylcarnitine | Acylcarnitines |
| C16 | Hexadecanoylcarnitine | Acylcarnitines |
| C16-OH | Hydroxyhexadecanoylcarnitine | Acylcarnitines |
| C16:1 | Hexadecenoylcarnitine | Acylcarnitines |
| C16:1-OH | Hydroxyhexadecenoylcarnitine | Acylcarnitines |
| C16:2 | Hexadecadienylcarnitine | Acylcarnitines |
| C16:2-OH | Hydroxyhexadecadienoylcarnitine | Acylcarnitines |
| C18 | Octadecanoylcarnitine | Acylcarnitines |
| C18:1 | Octadecenoylcarnitine | Acylcarnitines |
| C18:1-OH | Hydroxyoctadecenoylcarnitine | Acylcarnitines |
| C18:2 | Octadecadienylcarnitine | Acylcarnitines |
| Trigonelline | Trigonelline | Alkaloids |
| TMAO | Trimethylamine N-oxide | Amine Oxides |
| Ala | Alanine | Aminoacids |
| Arg | Arginine | Aminoacids |
| Asn | Asparagine | Aminoacids |
| Asp | Aspartic Acid | Aminoacids |
| Cys | Cysteine | Aminoacids |
| Gln | Glutamine | Aminoacids |
| Glu | Glutamic Acid | Aminoacids |
| Gly | Glycine | Aminoacids |
| His | Histidine | Aminoacids |
| Ile | Isoleucine | Aminoacids |
| Leu | Leucine | Aminoacids |
| Lys | Lysine | Aminoacids |
| Met | Methionine | Aminoacids |
| Phe | Phenylalanine | Aminoacids |
| Pro | Proline | Aminoacids |
| Ser | Serine | Aminoacids |
| Thr | Threonine | Aminoacids |
| Trp | Tryptophan | Aminoacids |
| Tyr | Tyrosine | Aminoacids |
| Val | Valine | Aminoacids |
| 1-Met-His | 1-Methylhistidine | Aminoacids Related |

| | | |
|---|---|---|
| 3-Met-His | 3-Methylhistidine | Aminoacids Related |
| 5-AVA | 5-Aminovaleric acid | Aminoacids Related |
| AABA | Alpha-Aminobutyric acid | Aminoacids Related |
| Ac-Orn | Acetylornithine | Aminoacids Related |
| ADMA | Asymmetric dimethylarginine | Aminoacids Related |
| alpha-AAA | alpha-Aminoadipic acid | Aminoacids Related |
| Anserine | L-Anserine | Aminoacids Related |
| BABA | beta-Aminobutyric acid | Aminoacids Related |
| Betaine | Betaine | Aminoacids Related |
| c4-OH-Pro | cis-4-Hydroxyproline | Aminoacids Related |
| Carnosine | Carnosine | Aminoacids Related |
| Cit | Citrulline | Aminoacids Related |
| Creatinine | Creatinine | Aminoacids Related |
| Cystine | Cystine | Aminoacids Related |
| DOPA | Dihydroxyphenylalanine | Aminoacids Related |
| HArg | Homoarginine | Aminoacids Related |
| HCys | Homocysteine | Aminoacids Related |
| Kynurenine | Kynurenine | Aminoacids Related |
| Met-SO | Methionine-Sulfoxide | Aminoacids Related |
| Nitro-Tyr | Nitrotyrosine | Aminoacids Related |
| Orn | Ornithine | Aminoacids Related |
| PAG | Phenylacetylglycine | Aminoacids Related |
| PheAlaBetaine | Phenylalanine betaine | Aminoacids Related |
| ProBetaine | Proline betaine | Aminoacids Related |
| Sarcosine | Sarcosine | Aminoacids Related |
| SDMA | Symmetric dimethylarginine | Aminoacids Related |
| t4-OH-Pro | trans-4-Hydroxyproline | Aminoacids Related |
| Taurine | Taurine | Aminoacids Related |
| TrpBetaine | Tryptophan betaine | Aminoacids Related |
| CA | Cholic Acid | Bile Acids |
| CDCA | Chenodeoxycholic acid | Bile Acids |
| DCA | Deoxycholic acid | Bile Acids |
| GCA | Glycocholic acid | Bile Acids |
| GCDCA | Glycochenodeoxycholic acid | Bile Acids |
| GDCA | Glycodeoxycholic acid | Bile Acids |
| GLCA | Glycolithocholic acid | Bile Acids |
| GLCAS | Glycolithocholic acid sulfate | Bile Acids |
| GUDCA | Glycoursodeoxycholic acid | Bile Acids |
| TCA | Taurocholic acid | Bile Acids |
| TCDCA | Taurochenodeoxycholic acid | Bile Acids |
| TDCA | Taurodeoxycholic acid | Bile Acids |
| TLCA | Taurolithocholic acid | Bile Acids |

| | | |
|---|---|---|
| TMCA | Tauro-muricholic acids | Bile Acids |
| beta-Ala | beta-Alanine | Biogenic Amines |
| Dopamine | Dopamine | Biogenic Amines |
| GABA | gamma-Aminobutyric acid | Biogenic Amines |
| Histamine | Histamine | Biogenic Amines |
| PEA | Phenylethylamine | Biogenic Amines |
| Putrescine | Putrescine | Biogenic Amines |
| Serotonin | Serotonin | Biogenic Amines |
| Spermidine | Spermidine | Biogenic Amines |
| Spermine | Spermine | Biogenic Amines |
| AconAcid | Aconitic acid | Carboxylic Acids |
| DiCA(12:0) | Dodecanedioic acid | Carboxylic Acids |
| DiCA(14:0) | Tetradecanedioic acid | Carboxylic Acids |
| HipAcid | Hippuric acid | Carboxylic Acids |
| Lac | Lactic acid | Carboxylic Acids |
| OH-GlutAcid | Hydroxyglutaric acid | Carboxylic Acids |
| Suc | Succinic acid | Carboxylic Acids |
| Cer(d16:1/18:0) | Ceramide(d16:1/18:0) | Ceramides |
| Cer(d16:1/20:0) | Ceramide(d16:1/20:0) | Ceramides |
| Cer(d16:1/22:0) | Ceramide(d16:1/22:0) | Ceramides |
| Cer(d16:1/23:0) | Ceramide(d16:1/23:0) | Ceramides |
| Cer(d16:1/24:0) | Ceramide(d16:1/24:0) | Ceramides |
| Cer(d18:1/14:0) | Ceramide(d18:1/14:0) | Ceramides |
| Cer(d18:1/16:0) | Ceramide(d18:1/16:0) | Ceramides |
| Cer(d18:1/18:0(OH)) | Ceramide(d18:1/18:0(OH)) | Ceramides |
| Cer(d18:1/18:0) | Ceramide(d18:1/18:0) | Ceramides |
| Cer(d18:1/18:1) | Ceramide(d18:1/18:1) | Ceramides |
| Cer(d18:1/20:0(OH)) | Ceramide(d18:1/20:0(OH)) | Ceramides |
| Cer(d18:1/20:0) | Ceramide(d18:1/20:0) | Ceramides |
| Cer(d18:1/22:0) | Ceramide(d18:1/22:0) | Ceramides |
| Cer(d18:1/23:0) | Ceramide(d18:1/23:0) | Ceramides |
| Cer(d18:1/24:0) | Ceramide(d18:1/24:0) | Ceramides |
| Cer(d18:1/24:1) | Ceramide(d18:1/24:1) | Ceramides |
| Cer(d18:1/25:0) | Ceramide(d18:1/25:0) | Ceramides |
| Cer(d18:1/26:0) | Ceramide(d18:1/26:0) | Ceramides |
| Cer(d18:1/26:1) | Ceramide(d18:1/26:1) | Ceramides |
| Cer(d18:2/14:0) | Ceramide(d18:2/14:0) | Ceramides |
| Cer(d18:2/16:0) | Ceramide(d18:2/16:0) | Ceramides |
| Cer(d18:2/18:0) | Ceramide(d18:2/18:0) | Ceramides |
| Cer(d18:2/18:1) | Ceramide(d18:2/18:1) | Ceramides |
| Cer(d18:2/20:0) | Ceramide(d18:2/20:0) | Ceramides |
| Cer(d18:2/22:0) | Ceramide(d18:2/22:0) | Ceramides |

| | | |
|---|---|---|
| Cer(d18:2/23:0) | Ceramide(d18:2/23:0) | Ceramides |
| Cer(d18:2/24:0) | Ceramide(d18:2/24:0) | Ceramides |
| Cer(d18:2/24:1) | Ceramide(d18:2/24:1) | Ceramides |
| CE(14:0) | Cholesteryl ester 14:0 | Cholesteryl Esters |
| CE(14:1) | Cholesteryl ester 14:1 | Cholesteryl Esters |
| CE(15:0) | Cholesteryl ester 15:0 | Cholesteryl Esters |
| CE(15:1) | Cholesteryl ester 15:1 | Cholesteryl Esters |
| CE(16:0) | Cholesteryl ester 16:0 | Cholesteryl Esters |
| CE(16:1) | Cholesteryl ester 16:1 | Cholesteryl Esters |
| CE(17:0) | Cholesteryl ester 17:0 | Cholesteryl Esters |
| CE(17:1) | Cholesteryl ester 17:1 | Cholesteryl Esters |
| CE(18:0) | Cholesteryl ester 18:0 | Cholesteryl Esters |
| CE(18:1) | Cholesteryl ester 18:1 | Cholesteryl Esters |
| CE(18:2) | Cholesteryl ester 18:2 | Cholesteryl Esters |
| CE(18:3) | Cholesteryl ester 18:3 | Cholesteryl Esters |
| CE(20:0) | Cholesteryl ester 20:0 | Cholesteryl Esters |
| CE(20:1) | Cholesteryl ester 20:1 | Cholesteryl Esters |
| CE(20:3) | Cholesteryl ester 20:3 | Cholesteryl Esters |
| CE(20:4) | Cholesteryl ester 20:4 | Cholesteryl Esters |
| CE(20:5) | Cholesteryl ester 20:5 | Cholesteryl Esters |
| CE(22:0) | Cholesteryl ester 22:0 | Cholesteryl Esters |
| CE(22:1) | Cholesteryl ester 22:1 | Cholesteryl Esters |
| CE(22:2) | Cholesteryl ester 22:2 | Cholesteryl Esters |
| CE(22:5) | Cholesteryl ester 22:5 | Cholesteryl Esters |
| CE(22:6) | Cholesteryl ester 22:6 | Cholesteryl Esters |
| p-Cresol-SO4 | p-Cresol sulfate | Cresols |
| DG(14:0_14:0) | Diacylglyceride(14:0_14:0) | Diacylglycerols |
| DG(14:0_18:1) | Diacylglyceride(14:0_18:1) | Diacylglycerols |
| DG(14:0_18:2) | Diacylglyceride(14:0_18:2) | Diacylglycerols |
| DG(14:0_20:0) | Diacylglyceride(14:0_20:0) | Diacylglycerols |
| DG(14:1_18:1) | Diacylglyceride(14:1_18:1) | Diacylglycerols |
| DG(14:1_20:2) | Diacylglyceride(14:1_20:2) | Diacylglycerols |
| DG(16:0_16:0) | Diacylglyceride(16:0_16:0) | Diacylglycerols |
| DG(16:0_16:1) | Diacylglyceride(16:0_16:1) | Diacylglycerols |
| DG(16:0_18:1) | Diacylglyceride(16:0_18:1) | Diacylglycerols |
| DG(16:0_18:2) | Diacylglyceride(16:0_18:2) | Diacylglycerols |
| DG(16:0_20:0) | Diacylglyceride(16:0_20:0) | Diacylglycerols |
| DG(16:0_20:3) | Diacylglyceride(16:0_20:3) | Diacylglycerols |
| DG(16:0_20:4) | Diacylglyceride(16:0_20:4) | Diacylglycerols |
| DG(16:1_18:0) | Diacylglyceride(16:1_18:0) | Diacylglycerols |
| DG(16:1_18:1) | Diacylglyceride(16:1_18:1) | Diacylglycerols |
| DG(16:1_18:2) | Diacylglyceride(16:1_18:2) | Diacylglycerols |

| | | |
|---|---|---|
| DG(16:1_20:0) | Diacylglyceride(16:1_20:0) | Diacylglycerols |
| DG(17:0_17:1) | Diacylglyceride(17:0_17:1) | Diacylglycerols |
| DG(17:0_18:1) | Diacylglyceride(17:0_18:1) | Diacylglycerols |
| DG(18:0_20:0) | Diacylglyceride(18:0_20:0) | Diacylglycerols |
| DG(18:0_20:4) | Diacylglyceride(18:0_20:4) | Diacylglycerols |
| DG(18:1_18:1) | Diacylglyceride(18:1_18:1) | Diacylglycerols |
| DG(18:1_18:2) | Diacylglyceride(18:1_18:2) | Diacylglycerols |
| DG(18:1_18:3) | Diacylglyceride(18:1_18:3) | Diacylglycerols |
| DG(18:1_18:4) | Diacylglyceride(18:1_18:4) | Diacylglycerols |
| DG(18:1_20:0) | Diacylglyceride(18:1_20:0) | Diacylglycerols |
| DG(18:1_20:1) | Diacylglyceride(18:1_20:1) | Diacylglycerols |
| DG(18:1_20:2) | Diacylglyceride(18:1_20:2) | Diacylglycerols |
| DG(18:1_20:3) | Diacylglyceride(18:1_20:3) | Diacylglycerols |
| DG(18:1_20:4) | Diacylglyceride(18:1_20:4) | Diacylglycerols |
| DG(18:1_22:5) | Diacylglyceride(18:1_22:5) | Diacylglycerols |
| DG(18:1_22:6) | Diacylglyceride(18:1_22:6) | Diacylglycerols |
| DG(18:2_18:2) | Diacylglyceride(18:2_18:2) | Diacylglycerols |
| DG(18:2_18:3) | Diacylglyceride(18:2_18:3) | Diacylglycerols |
| DG(18:2_18:4) | Diacylglyceride(18:2_18:4) | Diacylglycerols |
| DG(18:2_20:0) | Diacylglyceride(18:2_20:0) | Diacylglycerols |
| DG(18:2_20:4) | Diacylglyceride(18:2_20:4) | Diacylglycerols |
| DG(18:3_18:3) | Diacylglyceride(18:3_18:3) | Diacylglycerols |
| DG(18:3_20:2) | Diacylglyceride(18:3_20:2) | Diacylglycerols |
| DG(21:0_22:6) | Diacylglyceride(21:0_22:6) | Diacylglycerols |
| DG(22:1_22:2) | Diacylglyceride(22:1_22:2) | Diacylglycerols |
| DG-O(14:0_18:2) | Diacylglyceride O-(14:0_18:2) | Diacylglycerols |
| DG-O(16:0_18:1) | Diacylglyceride O-(16:0_18:1) | Diacylglycerols |
| DG-O(16:0_20:4) | Diacylglyceride O-(16:0_20:4) | Diacylglycerols |
| Cer(d18:0/18:0(OH)) | Dihydroceramide(d18:0/18:0(OH)) | Dihydroceramides |
| Cer(d18:0/18:0) | Dihydroceramide(d18:0/18:0) | Dihydroceramides |
| Cer(d18:0/20:0) | Dihydroceramide(d18:0/20:0) | Dihydroceramides |
| Cer(d18:0/22:0) | Dihydroceramide(d18:0/22:0) | Dihydroceramides |
| Cer(d18:0/24:0) | Dihydroceramide(d18:0/24:0) | Dihydroceramides |
| Cer(d18:0/24:1) | Dihydroceramide(d18:0/24:1) | Dihydroceramides |
| Cer(d18:0/26:1(OH)) | Dihydroceramide(d18:0/26:1(OH)) | Dihydroceramides |
| Cer(d18:0/26:1) | Dihydroceramide(d18:0/26:1) | Dihydroceramides |
| AA | Arachidonic acid | Fatty Acids |
| DHA | Docosahexaenoic acid | Fatty Acids |
| EPA | Eicosapentaenoic acid | Fatty Acids |
| FA(12:0) | Dodecanoic acid | Fatty Acids |
| FA(14:0) | Myristic acid | Fatty Acids |
| FA(16:0) | Palmitic acid | Fatty Acids |

| | | |
|---|---|---|
| FA(18:0) | Stearic acid | Fatty Acids |
| FA(18:1) | Octadecenoic acid | Fatty Acids |
| FA(18:2) | Octadecadienoate | Fatty Acids |
| FA(20:1) | Eicosenoic acid | Fatty Acids |
| FA(20:2) | Eicosadienoic acid | Fatty Acids |
| FA(20:3) | Eicosatrienoic acid | Fatty Acids |
| lysoPC a C14:0 | Lysophosphatidylcholine a C14:0 | Glycerophospholipids |
| lysoPC a C16:0 | Lysophosphatidylcholine a C16:0 | Glycerophospholipids |
| lysoPC a C16:1 | Lysophosphatidylcholine a C16:1 | Glycerophospholipids |
| lysoPC a C17:0 | Lysophosphatidylcholine a C17:0 | Glycerophospholipids |
| lysoPC a C18:0 | Lysophosphatidylcholine a C18:0 | Glycerophospholipids |
| lysoPC a C18:1 | Lysophosphatidylcholine a C18:1 | Glycerophospholipids |
| lysoPC a C18:2 | Lysophosphatidylcholine a C18:2 | Glycerophospholipids |
| lysoPC a C20:3 | Lysophosphatidylcholine a C20:3 | Glycerophospholipids |
| lysoPC a C20:4 | Lysophosphatidylcholine a C20:4 | Glycerophospholipids |
| lysoPC a C24:0 | Lysophosphatidylcholine a C24:0 | Glycerophospholipids |
| lysoPC a C26:0 | Lysophosphatidylcholine a C26:0 | Glycerophospholipids |
| lysoPC a C26:1 | Lysophosphatidylcholine a C26:1 | Glycerophospholipids |
| lysoPC a C28:0 | Lysophosphatidylcholine a C28:0 | Glycerophospholipids |
| lysoPC a C28:1 | Lysophosphatidylcholine a C28:1 | Glycerophospholipids |
| PC aa C24:0 | Phosphatidylcholine aa C24:0 | Glycerophospholipids |
| PC aa C26:0 | Phosphatidylcholine aa C26:0 | Glycerophospholipids |
| PC aa C28:1 | Phosphatidylcholine aa C28:1 | Glycerophospholipids |
| PC aa C30:0 | Phosphatidylcholine aa C30:0 | Glycerophospholipids |
| PC aa C30:2 | Phosphatidylcholine aa C30:2 | Glycerophospholipids |
| PC aa C32:0 | Phosphatidylcholine aa C32:0 | Glycerophospholipids |
| PC aa C32:1 | Phosphatidylcholine aa C32:1 | Glycerophospholipids |
| PC aa C32:2 | Phosphatidylcholine aa C32:2 | Glycerophospholipids |
| PC aa C32:3 | Phosphatidylcholine aa C32:3 | Glycerophospholipids |
| PC aa C34:1 | Phosphatidylcholine aa C34:1 | Glycerophospholipids |
| PC aa C34:2 | Phosphatidylcholine aa C34:2 | Glycerophospholipids |
| PC aa C34:3 | Phosphatidylcholine aa C34:3 | Glycerophospholipids |
| PC aa C34:4 | Phosphatidylcholine aa C34:4 | Glycerophospholipids |
| PC aa C36:0 | Phosphatidylcholine aa C36:0 | Glycerophospholipids |
| PC aa C36:1 | Phosphatidylcholine aa C36:1 | Glycerophospholipids |
| PC aa C36:2 | Phosphatidylcholine aa C36:2 | Glycerophospholipids |
| PC aa C36:3 | Phosphatidylcholine aa C36:3 | Glycerophospholipids |
| PC aa C36:4 | Phosphatidylcholine aa C36:4 | Glycerophospholipids |
| PC aa C36:5 | Phosphatidylcholine aa C36:5 | Glycerophospholipids |
| PC aa C36:6 | Phosphatidylcholine aa C36:6 | Glycerophospholipids |
| PC aa C38:0 | Phosphatidylcholine aa C38:0 | Glycerophospholipids |
| PC aa C38:1 | Phosphatidylcholine aa C38:1 | Glycerophospholipids |

| | | |
|---|---|---|
| PC aa C38:3 | Phosphatidylcholine aa C38:3 | Glycerophospholipids |
| PC aa C38:4 | Phosphatidylcholine aa C38:4 | Glycerophospholipids |
| PC aa C38:5 | Phosphatidylcholine aa C38:5 | Glycerophospholipids |
| PC aa C38:6 | Phosphatidylcholine aa C38:6 | Glycerophospholipids |
| PC aa C40:1 | Phosphatidylcholine aa C40:1 | Glycerophospholipids |
| PC aa C40:2 | Phosphatidylcholine aa C40:2 | Glycerophospholipids |
| PC aa C40:3 | Phosphatidylcholine aa C40:3 | Glycerophospholipids |
| PC aa C40:4 | Phosphatidylcholine aa C40:4 | Glycerophospholipids |
| PC aa C40:5 | Phosphatidylcholine aa C40:5 | Glycerophospholipids |
| PC aa C40:6 | Phosphatidylcholine aa C40:6 | Glycerophospholipids |
| PC aa C42:0 | Phosphatidylcholine aa C42:0 | Glycerophospholipids |
| PC aa C42:1 | Phosphatidylcholine aa C42:1 | Glycerophospholipids |
| PC aa C42:2 | Phosphatidylcholine aa C42:2 | Glycerophospholipids |
| PC aa C42:4 | Phosphatidylcholine aa C42:4 | Glycerophospholipids |
| PC aa C42:5 | Phosphatidylcholine aa C42:5 | Glycerophospholipids |
| PC aa C42:6 | Phosphatidylcholine aa C42:6 | Glycerophospholipids |
| PC ae C30:0 | Phosphatidylcholine ae C30:0 | Glycerophospholipids |
| PC ae C30:1 | Phosphatidylcholine ae C30:1 | Glycerophospholipids |
| PC ae C30:2 | Phosphatidylcholine ae C30:2 | Glycerophospholipids |
| PC ae C32:1 | Phosphatidylcholine ae C32:1 | Glycerophospholipids |
| PC ae C32:2 | Phosphatidylcholine ae C32:2 | Glycerophospholipids |
| PC ae C34:0 | Phosphatidylcholine ae C34:0 | Glycerophospholipids |
| PC ae C34:1 | Phosphatidylcholine ae C34:1 | Glycerophospholipids |
| PC ae C34:2 | Phosphatidylcholine ae C34:2 | Glycerophospholipids |
| PC ae C34:3 | Phosphatidylcholine ae C34:3 | Glycerophospholipids |
| PC ae C36:0 | Phosphatidylcholine ae C36:0 | Glycerophospholipids |
| PC ae C36:1 | Phosphatidylcholine ae C36:1 | Glycerophospholipids |
| PC ae C36:2 | Phosphatidylcholine ae C36:2 | Glycerophospholipids |
| PC ae C36:3 | Phosphatidylcholine ae C36:3 | Glycerophospholipids |
| PC ae C36:4 | Phosphatidylcholine ae C36:4 | Glycerophospholipids |
| PC ae C36:5 | Phosphatidylcholine ae C36:5 | Glycerophospholipids |
| PC ae C38:0 | Phosphatidylcholine ae C38:0 | Glycerophospholipids |
| PC ae C38:1 | Phosphatidylcholine ae C38:1 | Glycerophospholipids |
| PC ae C38:2 | Phosphatidylcholine ae C38:2 | Glycerophospholipids |
| PC ae C38:3 | Phosphatidylcholine ae C38:3 | Glycerophospholipids |
| PC ae C38:4 | Phosphatidylcholine ae C38:4 | Glycerophospholipids |
| PC ae C38:5 | Phosphatidylcholine ae C38:5 | Glycerophospholipids |
| PC ae C38:6 | Phosphatidylcholine ae C38:6 | Glycerophospholipids |
| PC ae C40:1 | Phosphatidylcholine ae C40:1 | Glycerophospholipids |
| PC ae C40:2 | Phosphatidylcholine ae C40:2 | Glycerophospholipids |
| PC ae C40:3 | Phosphatidylcholine ae C40:3 | Glycerophospholipids |
| PC ae C40:4 | Phosphatidylcholine ae C40:4 | Glycerophospholipids |

| | | |
|---|---|---|
| PC ae C40:5 | Phosphatidylcholine ae C40:5 | Glycerophospholipids |
| PC ae C40:6 | Phosphatidylcholine ae C40:6 | Glycerophospholipids |
| PC ae C42:0 | Phosphatidylcholine ae C42:0 | Glycerophospholipids |
| PC ae C42:1 | Phosphatidylcholine ae C42:1 | Glycerophospholipids |
| PC ae C42:2 | Phosphatidylcholine ae C42:2 | Glycerophospholipids |
| PC ae C42:3 | Phosphatidylcholine ae C42:3 | Glycerophospholipids |
| PC ae C42:4 | Phosphatidylcholine ae C42:4 | Glycerophospholipids |
| PC ae C42:5 | Phosphatidylcholine ae C42:5 | Glycerophospholipids |
| PC ae C44:3 | Phosphatidylcholine ae C44:3 | Glycerophospholipids |
| PC ae C44:4 | Phosphatidylcholine ae C44:4 | Glycerophospholipids |
| PC ae C44:5 | Phosphatidylcholine ae C44:5 | Glycerophospholipids |
| PC ae C44:6 | Phosphatidylcholine ae C44:6 | Glycerophospholipids |
| Hex2Cer(d18:1/14:0) | Dihexosylceramide(d18:1/14:0) | Glycosylceramides |
| Hex2Cer(d18:1/16:0) | Dihexosylceramide(d18:1/16:0) | Glycosylceramides |
| Hex2Cer(d18:1/18:0) | Dihexosylceramide(d18:1/18:0) | Glycosylceramides |
| Hex2Cer(d18:1/20:0) | Dihexosylceramide(d18:1/20:0) | Glycosylceramides |
| Hex2Cer(d18:1/22:0) | Dihexosylceramide(d18:1/22:0) | Glycosylceramides |
| Hex2Cer(d18:1/24:0) | Dihexosylceramide(d18:1/24:0) | Glycosylceramides |
| Hex2Cer(d18:1/24:1) | Dihexosylceramide(d18:1/24:1) | Glycosylceramides |
| Hex2Cer(d18:1/26:0) | Dihexosylceramide(d18:1/26:0) | Glycosylceramides |
| Hex2Cer(d18:1/26:1) | Dihexosylceramide(d18:1/26:1) | Glycosylceramides |
| Hex3Cer(d18:1/16:0) | Trihexosylceramide(d18:1/16:0) | Glycosylceramides |
| Hex3Cer(d18:1/18:0) | Trihexosylceramide(d18:1/18:0) | Glycosylceramides |
| Hex3Cer(d18:1/24:1) | Trihexosylceramide(d18:1/24:1) | Glycosylceramides |
| Hex3Cer(d18:1/26:1) | Trihexosylceramide(d18:1/26:1) | Glycosylceramides |
| Hex3Cer(d18:1_20:0) | Trihexosylceramide(d18:1_20:0) | Glycosylceramides |
| Hex3Cer(d18:1_22:0) | Trihexosylceramide(d18:1_22:0) | Glycosylceramides |
| HexCer(d16:1/22:0) | Hexosylceramide(d16:1/22:0) | Glycosylceramides |
| HexCer(d16:1/24:0) | Hexosylceramide(d16:1/24:0) | Glycosylceramides |
| HexCer(d18:1/14:0) | Hexosylceramide(d18:1/14:0) | Glycosylceramides |
| HexCer(d18:1/16:0) | Hexosylceramide(d18:1/16:0) | Glycosylceramides |
| HexCer(d18:1/18:0) | Hexosylceramide(d18:1/18:0) | Glycosylceramides |
| HexCer(d18:1/18:1) | Hexosylceramide(d18:1/18:1) | Glycosylceramides |
| HexCer(d18:1/20:0) | Hexosylceramide(d18:1/20:0) | Glycosylceramides |
| HexCer(d18:1/22:0) | Hexosylceramide(d18:1/22:0) | Glycosylceramides |
| HexCer(d18:1/23:0) | Hexosylceramide(d18:1/23:0) | Glycosylceramides |
| HexCer(d18:1/24:0) | Hexosylceramide(d18:1/24:0) | Glycosylceramides |
| HexCer(d18:1/24:1) | Hexosylceramide(d18:1/24:1) | Glycosylceramides |
| HexCer(d18:1/26:0) | Hexosylceramide(d18:1/26:0) | Glycosylceramides |
| HexCer(d18:1/26:1) | Hexosylceramide(d18:1/26:1) | Glycosylceramides |
| HexCer(d18:2/16:0) | Hexosylceramide(d18:2/16:0) | Glycosylceramides |
| HexCer(d18:2/18:0) | Hexosylceramide(d18:2/18:0) | Glycosylceramides |

| | | |
|---|---|---|
| HexCer(d18:2/20:0) | Hexosylceramide(d18:2/20:0) | Glycosylceramides |
| HexCer(d18:2/22:0) | Hexosylceramide(d18:2/22:0) | Glycosylceramides |
| HexCer(d18:2/23:0) | Hexosylceramide(d18:2/23:0) | Glycosylceramides |
| HexCer(d18:2/24:0) | Hexosylceramide(d18:2/24:0) | Glycosylceramides |
| AbsAcid | Abscisic acid | Hormones |
| Cortisol | Cortisol | Hormones |
| Cortisone | Cortisone | Hormones |
| DHEAS | Dehydroepiandrosterone sulfate | Hormones |
| 3-IAA | Indoleacetic acid | Indoles Derivatives |
| 3-IPA | Indolepropionic acid | Indoles Derivatives |
| Ind-SO4 | Indoxyl sulfate | Indoles Derivatives |
| Indole | Indole | Indoles Derivatives |
| Hypoxanthine | Hypoxanthine | Nucleobases Related |
| Xanthine | Xanthine | Nucleobases Related |
| SM (OH) C14:1 | Hydroxysphingomyelin C14:1 | Sphingolipids |
| SM (OH) C16:1 | Hydroxysphingomyelin C16:1 | Sphingolipids |
| SM (OH) C22:1 | Hydroxysphingomyelin C22:1 | Sphingolipids |
| SM (OH) C22:2 | Hydroxysphingomyelin C22:2 | Sphingolipids |
| SM (OH) C24:1 | Hydroxysphingomyelin C24:1 | Sphingolipids |
| SM C16:0 | Sphingomyelin C16:0 | Sphingolipids |
| SM C16:1 | Sphingomyelin C16:1 | Sphingolipids |
| SM C18:0 | Sphingomyelin C18:0 | Sphingolipids |
| SM C18:1 | Sphingomyelin C18:1 | Sphingolipids |
| SM C20:2 | Sphingomyelin C20:2 | Sphingolipids |
| SM C22:3 | Sphingomyelin C22:3 | Sphingolipids |
| SM C24:0 | Sphingomyelin C24:0 | Sphingolipids |
| SM C24:1 | Sphingomyelin C24:1 | Sphingolipids |
| SM C26:0 | Sphingomyelin C26:0 | Sphingolipids |
| SM C26:1 | Sphingomyelin C26:1 | Sphingolipids |
| H1 | Hexose | Sugars |
| TG(14:0_32:2) | Triacylglyceride(14:0_32:2) | Triacylglycerols |
| TG(14:0_34:0) | Triacylglyceride(14:0_34:0) | Triacylglycerols |
| TG(14:0_34:1) | Triacylglyceride(14:0_34:1) | Triacylglycerols |
| TG(14:0_34:2) | Triacylglyceride(14:0_34:2) | Triacylglycerols |
| TG(14:0_34:3) | Triacylglyceride(14:0_34:3) | Triacylglycerols |
| TG(14:0_35:1) | Triacylglyceride(14:0_35:1) | Triacylglycerols |
| TG(14:0_35:2) | Triacylglyceride(14:0_35:2) | Triacylglycerols |
| TG(14:0_36:1) | Triacylglyceride(14:0_36:1) | Triacylglycerols |
| TG(14:0_36:2) | Triacylglyceride(14:0_36:2) | Triacylglycerols |
| TG(14:0_36:3) | Triacylglyceride(14:0_36:3) | Triacylglycerols |
| TG(14:0_36:4) | Triacylglyceride(14:0_36:4) | Triacylglycerols |
| TG(14:0_38:4) | Triacylglyceride(14:0_38:4) | Triacylglycerols |

| | | |
|---|---|---|
| TG(14:0_38:5) | Triacylglyceride(14:0_38:5) | Triacylglycerols |
| TG(14:0_39:3) | Triacylglyceride(14:0_39:3) | Triacylglycerols |
| TG(16:0_28:1) | Triacylglyceride(16:0_28:1) | Triacylglycerols |
| TG(16:0_28:2) | Triacylglyceride(16:0_28:2) | Triacylglycerols |
| TG(16:0_30:2) | Triacylglyceride(16:0_30:2) | Triacylglycerols |
| TG(16:0_32:0) | Triacylglyceride(16:0_32:0) | Triacylglycerols |
| TG(16:0_32:1) | Triacylglyceride(16:0_32:1) | Triacylglycerols |
| TG(16:0_32:2) | Triacylglyceride(16:0_32:2) | Triacylglycerols |
| TG(16:0_32:3) | Triacylglyceride(16:0_32:3) | Triacylglycerols |
| TG(16:0_33:1) | Triacylglyceride(16:0_33:1) | Triacylglycerols |
| TG(16:0_33:2) | Triacylglyceride(16:0_33:2) | Triacylglycerols |
| TG(16:0_34:0) | Triacylglyceride(16:0_34:0) | Triacylglycerols |
| TG(16:0_34:1) | Triacylglyceride(16:0_34:1) | Triacylglycerols |
| TG(16:0_34:2) | Triacylglyceride(16:0_34:2) | Triacylglycerols |
| TG(16:0_34:3) | Triacylglyceride(16:0_34:3) | Triacylglycerols |
| TG(16:0_34:4) | Triacylglyceride(16:0_34:4) | Triacylglycerols |
| TG(16:0_35:1) | Triacylglyceride(16:0_35:1) | Triacylglycerols |
| TG(16:0_35:2) | Triacylglyceride(16:0_35:2) | Triacylglycerols |
| TG(16:0_35:3) | Triacylglyceride(16:0_35:3) | Triacylglycerols |
| TG(16:0_36:2) | Triacylglyceride(16:0_36:2) | Triacylglycerols |
| TG(16:0_36:3) | Triacylglyceride(16:0_36:3) | Triacylglycerols |
| TG(16:0_36:4) | Triacylglyceride(16:0_36:4) | Triacylglycerols |
| TG(16:0_36:5) | Triacylglyceride(16:0_36:5) | Triacylglycerols |
| TG(16:0_36:6) | Triacylglyceride(16:0_36:6) | Triacylglycerols |
| TG(16:0_37:3) | Triacylglyceride(16:0_37:3) | Triacylglycerols |
| TG(16:0_38:1) | Triacylglyceride(16:0_38:1) | Triacylglycerols |
| TG(16:0_38:2) | Triacylglyceride(16:0_38:2) | Triacylglycerols |
| TG(16:0_38:3) | Triacylglyceride(16:0_38:3) | Triacylglycerols |
| TG(16:0_38:4) | Triacylglyceride(16:0_38:4) | Triacylglycerols |
| TG(16:0_38:5) | Triacylglyceride(16:0_38:5) | Triacylglycerols |
| TG(16:0_38:6) | Triacylglyceride(16:0_38:6) | Triacylglycerols |
| TG(16:0_38:7) | Triacylglyceride(16:0_38:7) | Triacylglycerols |
| TG(16:0_40:6) | Triacylglyceride(16:0_40:6) | Triacylglycerols |
| TG(16:0_40:7) | Triacylglyceride(16:0_40:7) | Triacylglycerols |
| TG(16:0_40:8) | Triacylglyceride(16:0_40:8) | Triacylglycerols |
| TG(16:1_28:0) | Triacylglyceride(16:1_28:0) | Triacylglycerols |
| TG(16:1_30:1) | Triacylglyceride(16:1_30:1) | Triacylglycerols |
| TG(16:1_32:0) | Triacylglyceride(16:1_32:0) | Triacylglycerols |
| TG(16:1_32:1) | Triacylglyceride(16:1_32:1) | Triacylglycerols |
| TG(16:1_32:2) | Triacylglyceride(16:1_32:2) | Triacylglycerols |
| TG(16:1_33:1) | Triacylglyceride(16:1_33:1) | Triacylglycerols |
| TG(16:1_34:0) | Triacylglyceride(16:1_34:0) | Triacylglycerols |

| | | |
|---|---|---|
| TG(16:1_34:1) | Triacylglyceride(16:1_34:1) | Triacylglycerols |
| TG(16:1_34:2) | Triacylglyceride(16:1_34:2) | Triacylglycerols |
| TG(16:1_34:3) | Triacylglyceride(16:1_34:3) | Triacylglycerols |
| TG(16:1_36:1) | Triacylglyceride(16:1_36:1) | Triacylglycerols |
| TG(16:1_36:2) | Triacylglyceride(16:1_36:2) | Triacylglycerols |
| TG(16:1_36:3) | Triacylglyceride(16:1_36:3) | Triacylglycerols |
| TG(16:1_36:4) | Triacylglyceride(16:1_36:4) | Triacylglycerols |
| TG(16:1_36:5) | Triacylglyceride(16:1_36:5) | Triacylglycerols |
| TG(16:1_38:3) | Triacylglyceride(16:1_38:3) | Triacylglycerols |
| TG(16:1_38:4) | Triacylglyceride(16:1_38:4) | Triacylglycerols |
| TG(16:1_38:5) | Triacylglyceride(16:1_38:5) | Triacylglycerols |
| TG(17:0_32:1) | Triacylglyceride(17:0_32:1) | Triacylglycerols |
| TG(17:0_34:1) | Triacylglyceride(17:0_34:1) | Triacylglycerols |
| TG(17:0_34:2) | Triacylglyceride(17:0_34:2) | Triacylglycerols |
| TG(17:0_34:3) | Triacylglyceride(17:0_34:3) | Triacylglycerols |
| TG(17:0_36:3) | Triacylglyceride(17:0_36:3) | Triacylglycerols |
| TG(17:0_36:4) | Triacylglyceride(17:0_36:4) | Triacylglycerols |
| TG(17:1_32:1) | Triacylglyceride(17:1_32:1) | Triacylglycerols |
| TG(17:1_34:1) | Triacylglyceride(17:1_34:1) | Triacylglycerols |
| TG(17:1_34:2) | Triacylglyceride(17:1_34:2) | Triacylglycerols |
| TG(17:1_34:3) | Triacylglyceride(17:1_34:3) | Triacylglycerols |
| TG(17:1_36:3) | Triacylglyceride(17:1_36:3) | Triacylglycerols |
| TG(17:1_36:4) | Triacylglyceride(17:1_36:4) | Triacylglycerols |
| TG(17:1_36:5) | Triacylglyceride(17:1_36:5) | Triacylglycerols |
| TG(17:1_38:5) | Triacylglyceride(17:1_38:5) | Triacylglycerols |
| TG(17:1_38:6) | Triacylglyceride(17:1_38:6) | Triacylglycerols |
| TG(17:1_38:7) | Triacylglyceride(17:1_38:7) | Triacylglycerols |
| TG(17:2_34:2) | Triacylglyceride(17:2_34:2) | Triacylglycerols |
| TG(17:2_34:3) | Triacylglyceride(17:2_34:3) | Triacylglycerols |
| TG(17:2_36:2) | Triacylglyceride(17:2_36:2) | Triacylglycerols |
| TG(17:2_36:3) | Triacylglyceride(17:2_36:3) | Triacylglycerols |
| TG(17:2_36:4) | Triacylglyceride(17:2_36:4) | Triacylglycerols |
| TG(17:2_38:5) | Triacylglyceride(17:2_38:5) | Triacylglycerols |
| TG(17:2_38:6) | Triacylglyceride(17:2_38:6) | Triacylglycerols |
| TG(17:2_38:7) | Triacylglyceride(17:2_38:7) | Triacylglycerols |
| TG(18:0_30:0) | Triacylglyceride(18:0_30:0) | Triacylglycerols |
| TG(18:0_30:1) | Triacylglyceride(18:0_30:1) | Triacylglycerols |
| TG(18:0_32:0) | Triacylglyceride(18:0_32:0) | Triacylglycerols |
| TG(18:0_32:1) | Triacylglyceride(18:0_32:1) | Triacylglycerols |
| TG(18:0_32:2) | Triacylglyceride(18:0_32:2) | Triacylglycerols |
| TG(18:0_34:2) | Triacylglyceride(18:0_34:2) | Triacylglycerols |
| TG(18:0_34:3) | Triacylglyceride(18:0_34:3) | Triacylglycerols |

| | | |
|---|---|---|
| TG(18:0_36:1) | Triacylglyceride(18:0_36:1) | Triacylglycerols |
| TG(18:0_36:2) | Triacylglyceride(18:0_36:2) | Triacylglycerols |
| TG(18:0_36:3) | Triacylglyceride(18:0_36:3) | Triacylglycerols |
| TG(18:0_36:4) | Triacylglyceride(18:0_36:4) | Triacylglycerols |
| TG(18:0_36:5) | Triacylglyceride(18:0_36:5) | Triacylglycerols |
| TG(18:0_38:6) | Triacylglyceride(18:0_38:6) | Triacylglycerols |
| TG(18:0_38:7) | Triacylglyceride(18:0_38:7) | Triacylglycerols |
| TG(18:1_26:0) | Triacylglyceride(18:1_26:0) | Triacylglycerols |
| TG(18:1_28:1) | Triacylglyceride(18:1_28:1) | Triacylglycerols |
| TG(18:1_30:0) | Triacylglyceride(18:1_30:0) | Triacylglycerols |
| TG(18:1_30:1) | Triacylglyceride(18:1_30:1) | Triacylglycerols |
| TG(18:1_30:2) | Triacylglyceride(18:1_30:2) | Triacylglycerols |
| TG(18:1_31:0) | Triacylglyceride(18:1_31:0) | Triacylglycerols |
| TG(18:1_32:0) | Triacylglyceride(18:1_32:0) | Triacylglycerols |
| TG(18:1_32:1) | Triacylglyceride(18:1_32:1) | Triacylglycerols |
| TG(18:1_32:2) | Triacylglyceride(18:1_32:2) | Triacylglycerols |
| TG(18:1_32:3) | Triacylglyceride(18:1_32:3) | Triacylglycerols |
| TG(18:1_33:0) | Triacylglyceride(18:1_33:0) | Triacylglycerols |
| TG(18:1_33:1) | Triacylglyceride(18:1_33:1) | Triacylglycerols |
| TG(18:1_33:2) | Triacylglyceride(18:1_33:2) | Triacylglycerols |
| TG(18:1_33:3) | Triacylglyceride(18:1_33:3) | Triacylglycerols |
| TG(18:1_34:1) | Triacylglyceride(18:1_34:1) | Triacylglycerols |
| TG(18:1_34:2) | Triacylglyceride(18:1_34:2) | Triacylglycerols |
| TG(18:1_34:3) | Triacylglyceride(18:1_34:3) | Triacylglycerols |
| TG(18:1_34:4) | Triacylglyceride(18:1_34:4) | Triacylglycerols |
| TG(18:1_35:2) | Triacylglyceride(18:1_35:2) | Triacylglycerols |
| TG(18:1_35:3) | Triacylglyceride(18:1_35:3) | Triacylglycerols |
| TG(18:1_36:0) | Triacylglyceride(18:1_36:0) | Triacylglycerols |
| TG(18:1_36:1) | Triacylglyceride(18:1_36:1) | Triacylglycerols |
| TG(18:1_36:2) | Triacylglyceride(18:1_36:2) | Triacylglycerols |
| TG(18:1_36:3) | Triacylglyceride(18:1_36:3) | Triacylglycerols |
| TG(18:1_36:4) | Triacylglyceride(18:1_36:4) | Triacylglycerols |
| TG(18:1_36:5) | Triacylglyceride(18:1_36:5) | Triacylglycerols |
| TG(18:1_36:6) | Triacylglyceride(18:1_36:6) | Triacylglycerols |
| TG(18:1_38:5) | Triacylglyceride(18:1_38:5) | Triacylglycerols |
| TG(18:1_38:6) | Triacylglyceride(18:1_38:6) | Triacylglycerols |
| TG(18:1_38:7) | Triacylglyceride(18:1_38:7) | Triacylglycerols |
| TG(18:2_28:0) | Triacylglyceride(18:2_28:0) | Triacylglycerols |
| TG(18:2_30:0) | Triacylglyceride(18:2_30:0) | Triacylglycerols |
| TG(18:2_30:1) | Triacylglyceride(18:2_30:1) | Triacylglycerols |
| TG(18:2_31:0) | Triacylglyceride(18:2_31:0) | Triacylglycerols |
| TG(18:2_32:0) | Triacylglyceride(18:2_32:0) | Triacylglycerols |

| | | |
|---|---|---|
| TG(18:2_32:1) | Triacylglyceride(18:2_32:1) | Triacylglycerols |
| TG(18:2_32:2) | Triacylglyceride(18:2_32:2) | Triacylglycerols |
| TG(18:2_33:0) | Triacylglyceride(18:2_33:0) | Triacylglycerols |
| TG(18:2_33:1) | Triacylglyceride(18:2_33:1) | Triacylglycerols |
| TG(18:2_33:2) | Triacylglyceride(18:2_33:2) | Triacylglycerols |
| TG(18:2_34:0) | Triacylglyceride(18:2_34:0) | Triacylglycerols |
| TG(18:2_34:1) | Triacylglyceride(18:2_34:1) | Triacylglycerols |
| TG(18:2_34:2) | Triacylglyceride(18:2_34:2) | Triacylglycerols |
| TG(18:2_34:3) | Triacylglyceride(18:2_34:3) | Triacylglycerols |
| TG(18:2_34:4) | Triacylglyceride(18:2_34:4) | Triacylglycerols |
| TG(18:2_35:1) | Triacylglyceride(18:2_35:1) | Triacylglycerols |
| TG(18:2_35:2) | Triacylglyceride(18:2_35:2) | Triacylglycerols |
| TG(18:2_35:3) | Triacylglyceride(18:2_35:3) | Triacylglycerols |
| TG(18:2_36:0) | Triacylglyceride(18:2_36:0) | Triacylglycerols |
| TG(18:2_36:1) | Triacylglyceride(18:2_36:1) | Triacylglycerols |
| TG(18:2_36:2) | Triacylglyceride(18:2_36:2) | Triacylglycerols |
| TG(18:2_36:3) | Triacylglyceride(18:2_36:3) | Triacylglycerols |
| TG(18:2_36:4) | Triacylglyceride(18:2_36:4) | Triacylglycerols |
| TG(18:2_36:5) | Triacylglyceride(18:2_36:5) | Triacylglycerols |
| TG(18:2_38:4) | Triacylglyceride(18:2_38:4) | Triacylglycerols |
| TG(18:2_38:5) | Triacylglyceride(18:2_38:5) | Triacylglycerols |
| TG(18:2_38:6) | Triacylglyceride(18:2_38:6) | Triacylglycerols |
| TG(18:3_30:0) | Triacylglyceride(18:3_30:0) | Triacylglycerols |
| TG(18:3_32:0) | Triacylglyceride(18:3_32:0) | Triacylglycerols |
| TG(18:3_32:1) | Triacylglyceride(18:3_32:1) | Triacylglycerols |
| TG(18:3_33:2) | Triacylglyceride(18:3_33:2) | Triacylglycerols |
| TG(18:3_34:0) | Triacylglyceride(18:3_34:0) | Triacylglycerols |
| TG(18:3_34:1) | Triacylglyceride(18:3_34:1) | Triacylglycerols |
| TG(18:3_34:2) | Triacylglyceride(18:3_34:2) | Triacylglycerols |
| TG(18:3_34:3) | Triacylglyceride(18:3_34:3) | Triacylglycerols |
| TG(18:3_35:2) | Triacylglyceride(18:3_35:2) | Triacylglycerols |
| TG(18:3_36:1) | Triacylglyceride(18:3_36:1) | Triacylglycerols |
| TG(18:3_36:2) | Triacylglyceride(18:3_36:2) | Triacylglycerols |
| TG(18:3_36:3) | Triacylglyceride(18:3_36:3) | Triacylglycerols |
| TG(18:3_36:4) | Triacylglyceride(18:3_36:4) | Triacylglycerols |
| TG(18:3_38:5) | Triacylglyceride(18:3_38:5) | Triacylglycerols |
| TG(18:3_38:6) | Triacylglyceride(18:3_38:6) | Triacylglycerols |
| TG(20:0_32:3) | Triacylglyceride(20:0_32:3) | Triacylglycerols |
| TG(20:0_32:4) | Triacylglyceride(20:0_32:4) | Triacylglycerols |
| TG(20:0_34:1) | Triacylglyceride(20:0_34:1) | Triacylglycerols |
| TG(20:1_24:3) | Triacylglyceride(20:1_24:3) | Triacylglycerols |
| TG(20:1_26:1) | Triacylglyceride(20:1_26:1) | Triacylglycerols |

| | | |
|---|---|---|
| TG(20:1_30:1) | Triacylglyceride(20:1_30:1) | Triacylglycerols |
| TG(20:1_31:0) | Triacylglyceride(20:1_31:0) | Triacylglycerols |
| TG(20:1_32:1) | Triacylglyceride(20:1_32:1) | Triacylglycerols |
| TG(20:1_32:2) | Triacylglyceride(20:1_32:2) | Triacylglycerols |
| TG(20:1_32:3) | Triacylglyceride(20:1_32:3) | Triacylglycerols |
| TG(20:1_34:0) | Triacylglyceride(20:1_34:0) | Triacylglycerols |
| TG(20:1_34:1) | Triacylglyceride(20:1_34:1) | Triacylglycerols |
| TG(20:1_34:2) | Triacylglyceride(20:1_34:2) | Triacylglycerols |
| TG(20:1_34:3) | Triacylglyceride(20:1_34:3) | Triacylglycerols |
| TG(20:2_32:0) | Triacylglyceride(20:2_32:0) | Triacylglycerols |
| TG(20:2_32:1) | Triacylglyceride(20:2_32:1) | Triacylglycerols |
| TG(20:2_34:1) | Triacylglyceride(20:2_34:1) | Triacylglycerols |
| TG(20:2_34:2) | Triacylglyceride(20:2_34:2) | Triacylglycerols |
| TG(20:2_34:3) | Triacylglyceride(20:2_34:3) | Triacylglycerols |
| TG(20:2_34:4) | Triacylglyceride(20:2_34:4) | Triacylglycerols |
| TG(20:2_36:5) | Triacylglyceride(20:2_36:5) | Triacylglycerols |
| TG(20:3_32:0) | Triacylglyceride(20:3_32:0) | Triacylglycerols |
| TG(20:3_32:1) | Triacylglyceride(20:3_32:1) | Triacylglycerols |
| TG(20:3_32:2) | Triacylglyceride(20:3_32:2) | Triacylglycerols |
| TG(20:3_34:0) | Triacylglyceride(20:3_34:0) | Triacylglycerols |
| TG(20:3_34:1) | Triacylglyceride(20:3_34:1) | Triacylglycerols |
| TG(20:3_34:2) | Triacylglyceride(20:3_34:2) | Triacylglycerols |
| TG(20:3_34:3) | Triacylglyceride(20:3_34:3) | Triacylglycerols |
| TG(20:3_36:3) | Triacylglyceride(20:3_36:3) | Triacylglycerols |
| TG(20:3_36:4) | Triacylglyceride(20:3_36:4) | Triacylglycerols |
| TG(20:3_36:5) | Triacylglyceride(20:3_36:5) | Triacylglycerols |
| TG(20:4_30:0) | Triacylglyceride(20:4_30:0) | Triacylglycerols |
| TG(20:4_32:0) | Triacylglyceride(20:4_32:0) | Triacylglycerols |
| TG(20:4_32:1) | Triacylglyceride(20:4_32:1) | Triacylglycerols |
| TG(20:4_32:2) | Triacylglyceride(20:4_32:2) | Triacylglycerols |
| TG(20:4_33:2) | Triacylglyceride(20:4_33:2) | Triacylglycerols |
| TG(20:4_34:0) | Triacylglyceride(20:4_34:0) | Triacylglycerols |
| TG(20:4_34:1) | Triacylglyceride(20:4_34:1) | Triacylglycerols |
| TG(20:4_34:2) | Triacylglyceride(20:4_34:2) | Triacylglycerols |
| TG(20:4_34:3) | Triacylglyceride(20:4_34:3) | Triacylglycerols |
| TG(20:4_35:3) | Triacylglyceride(20:4_35:3) | Triacylglycerols |
| TG(20:4_36:2) | Triacylglyceride(20:4_36:2) | Triacylglycerols |
| TG(20:4_36:3) | Triacylglyceride(20:4_36:3) | Triacylglycerols |
| TG(20:4_36:4) | Triacylglyceride(20:4_36:4) | Triacylglycerols |
| TG(20:4_36:5) | Triacylglyceride(20:4_36:5) | Triacylglycerols |
| TG(20:5_34:0) | Triacylglyceride(20:5_34:0) | Triacylglycerols |
| TG(20:5_34:1) | Triacylglyceride(20:5_34:1) | Triacylglycerols |

| | | |
|---|---|---|
| TG(20:5_34:2) | Triacylglyceride(20:5_34:2) | Triacylglycerols |
| TG(20:5_36:2) | Triacylglyceride(20:5_36:2) | Triacylglycerols |
| TG(20:5_36:3) | Triacylglyceride(20:5_36:3) | Triacylglycerols |
| TG(22:0_32:4) | Triacylglyceride(22:0_32:4) | Triacylglycerols |
| TG(22:1_32:5) | Triacylglyceride(22:1_32:5) | Triacylglycerols |
| TG(22:2_32:4) | Triacylglyceride(22:2_32:4) | Triacylglycerols |
| TG(22:3_30:2) | Triacylglyceride(22:3_30:2) | Triacylglycerols |
| TG(22:4_32:0) | Triacylglyceride(22:4_32:0) | Triacylglycerols |
| TG(22:4_32:2) | Triacylglyceride(22:4_32:2) | Triacylglycerols |
| TG(22:4_34:2) | Triacylglyceride(22:4_34:2) | Triacylglycerols |
| TG(22:5_32:0) | Triacylglyceride(22:5_32:0) | Triacylglycerols |
| TG(22:5_32:1) | Triacylglyceride(22:5_32:1) | Triacylglycerols |
| TG(22:5_34:1) | Triacylglyceride(22:5_34:1) | Triacylglycerols |
| TG(22:5_34:2) | Triacylglyceride(22:5_34:2) | Triacylglycerols |
| TG(22:5_34:3) | Triacylglyceride(22:5_34:3) | Triacylglycerols |
| TG(22:6_32:0) | Triacylglyceride(22:6_32:0) | Triacylglycerols |
| TG(22:6_32:1) | Triacylglyceride(22:6_32:1) | Triacylglycerols |
| TG(22:6_34:1) | Triacylglyceride(22:6_34:1) | Triacylglycerols |
| TG(22:6_34:2) | Triacylglyceride(22:6_34:2) | Triacylglycerols |
| TG(22:6_34:3) | Triacylglyceride(22:6_34:3) | Triacylglycerols |
| Choline | Choline | Vitamins & Cofactors |
